# Introspection confidence predicts EEG decoding of self-generated thoughts and meta-awareness

**DOI:** 10.1101/2021.03.12.435068

**Authors:** Naya Polychroni, Maria Herrojo Ruiz, Devin B. Terhune

## Abstract

The neurophysiological bases of mind wandering (MW) – an experiential state wherein attention is disengaged from the external environment in favour of internal thoughts, and state meta-awareness are poorly understood. In parallel, the relationship between introspection confidence in experiential state judgements and neural representations remains unclear. Here, we recorded EEG whilst participants completed a listening task within which they made experiential state judgments and rated their confidence. Alpha power was reliably greater during MW episodes, with unaware MW further associated with greater delta and theta power. Multivariate pattern classification analysis revealed that MW, and meta-awareness can be decoded from the distribution of power in these three frequency bands. Critically, we show that individual decoding accuracies positively correlate with introspection confidence. Our results reaffirm the role of alpha oscillations in MW, implicate lower frequencies in meta-awareness, and are consistent with the proposal that introspection confidence indexes neurophysiological discriminability of representational states.

## 1 Introduction

Our brains are constantly bombarded by dynamic sensory input, yet we frequently shift away from the external environment towards thoughts, emotions and images that do not emerge from ongoing perceptual processes, are self-generated and unrelated to one’s current task (*mind wandering*; (Antrobus et al., 1970; Christoff et al., 2009; Smallwood & Schooler, 2015). Research suggests that mind wandering (MW) occurs in ∼30-50% of our waking hours (Killingsworth & Gilbert, 2010; McVay et al., 2009), has deleterious effects on sensory and cognitive processing with corresponding reductions in event-related potentials (ERP) in response to external stimuli (Smallwood et al. 2008; Barron et al. 2011; Kam et al. 2011; Baird et al. 2014). These effects produce concomitant negative effects in a variety of tasks from driving (Geden et al., 2018) to reading (Feng et al., 2013; Hollis & Was, 2016; Schooler et al., 2004).

Converging findings implicate default mode network (DMN; Fox et al., 2005; Raichle et al., 2001) in mind wandering (Christoff et al., 2009) and its associated cognitive operations such as self-related processing (Fingelkurts & Fingelkurts, 2011), autobiographical memory, theory of mind, and future planning (Andrews-Hanna, 2012; Andrews-Hanna et al., 2014; Spreng & Grady, 2010). Mind wandering episodes can occur with (*tuning-out*) and without (*zoning-out*) meta-awareness (Schooler et al. 2004, 2011; Smallwood, McSpadden, et al. 2007) with the latter suggested to reflect a more pronounced form of mind wandering characterized by poorer performance (Smallwood, McSpadden, and Schooler 2007) and greater recruitment of DMN and executive control network (Christoff et al., 2009).

Despite advances in its network architecture, the patterns of oscillatory activity underpinning mind wandering are poorly understood (Martinon et al., 2019). Multiple studies indicate that mind wandering states (Boudewyn & Carter, 2018; Compton et al., 2019; Groot et al., 2021; Macdonald et al., 2011), and particularly zoning-out (unaware mind wandering) (Boudewyn and Carter 2018), are characterized by elevated alpha (∼8-12Hz) power. Alpha oscillations are suggested to support inhibition-related processes (Klimesch, 2012; Palva & Palva, 2007), and attentional suppression (Foxe & Snyder, 2011) and are further implicated during working memory and mental imagery tasks (Von Stein & Sarnthein, 2000), internally-oriented brain states (Cooper et al., 2003; Hanslmayr et al., 2011), and inner speech (Villena-González et al., 2016), all of which figure prominently in the experience of mind wandering. However, at least two studies failed to replicate these effects (Braboszcz & Delorme, 2011; van Son et al., 2019) and observed greater delta (∼2-3Hz) and theta (∼4-7Hz) power during mind wandering. Activity in these frequency bands may reflect lapses in sustained attention, redirection of attention from external stimuli to internal representations and, or the maintenance of these representations in awareness. For example, slow-wave brain oscillations are typically associated with decreased sustained task-related attention (Klimesch, 1999). Delta frequency contributions have also been shown during increased focus on internal processing and pertinent inhibition of interference (Harmony, 2013) whereas theta activity has been consistently shown to relate to maintenance of information in working memory (Klimesch, 1999; Mitchell et al., 2008). Discrepancies in the observed association between alpha and mind wandering are plausibly attributed to the tasks and methods used in the aforementioned studies (Braboszcz & Delorme, 2011; van Son et al., 2019). That is, lower alpha activity during mind wandering episodes in some studies might be due to the concurrent task (breath-counting) involving internally-focused attention and counting (Palva et al. 2005; Sauseng et al. 2005). In parallel, it is difficult to compare findings between self-reports that are prompted (probe-caught) and the above studies due to the latter using self-caught measures (participants are asked to indicate when they catch themselves mind wandering), which likely capture shifts from internal to external focus that probably involve different mechanisms to the occurrence of mind wandering.

Measuring the neurophysiology of self-generated thoughts requires experience sampling methods in which participants report on aspects of their experience, thereby affording a prominent role to introspective abilities in the assessment of mind wandering (Smallwood & Schooler, 2006). A neglected feature of these abilities within the context of mind wandering is confidence in these introspective reports. Emerging evidence suggests that confidence reflects variability in access to experiential states (Fleming & Lau 2014; Seli et al. 2015) and thus is likely to be highly informative in elucidating variability in the phenomenology and neurophysiology of mind wandering episodes. Confidence in perceptual judgements (Fleming & Lau, 2014) positively correlates with decision accuracy (Gherman & Philiastides, 2018; Kunimoto et al., 2001; Morgan et al., 1997), and reliably tracks ERP dynamics related to error (Boldt & Yeung, 2015) and sensory processing (e.g. Zakrzewski et al. 2019). Confidence in mind wandering reports has to date been neglected but preliminary work has shown that it varies greatly within and between individuals and moderates the relationship between response time variability and self-reports of mind wandering (Seli et al. 2015) (but see Meier 2018). Nevertheless, the neurophysiology of these effects is unknown.

One possibility is that if confidence reflects superior access to experiential states, high confidence would be associated with more clearly dissociable neural representations. Multivariate pattern classification analysis (MVPC) has been successfully used to identify the mapping between distributed patterns of neural activity and corresponding mental states (Bae & Luck, 2018; Haxby et al., 2001; Haynes & Rees, 2006; Jin et al., 2019; Mittner et al., 2014). An advantage of MVPC is that it allows researchers to assess whether shared information across multiple features (e.g., channels, frequency, time points) encodes class-related information (Haxby et al. 2001). Using this method, recent research has revealed associations between decoding accuracy and individual differences in perceptual discrimination (Kim et al. 2015) and intra-individual variability in confidence (Weaver et al., 2019). The extent to which experiential states can be decoded, reflecting multivariate dissimilarity of neural representations, may thus underlie confidence in the corresponding mental representations.

The present study investigated the oscillatory dynamics of experiential states using an ecological task lacking performance indicators (Smallwood et al., 2009) in order to examine the neurophysiological basis of mind wandering, dissociate meta-awareness of mind wandering (henceforth state meta-awareness) and investigate the neurophysiological implications of participants’ confidence in self-reports. During concurrent EEG recording, participants listened to an audiobook and were intermittently probed regarding their experiential state and state meta-awareness, and rated their confidence in both judgements. We expected that mind wandering would be characterized by elevated alpha power (Compton et al. 2019) whereas unaware mind wandering would additionally be associated with differential power in slow oscillatory bands (delta, theta).

Motivated by previous research showing that joint activity patterns across different features (e.g. frequency bands) can be more informative of mental representations than univariate information (Allefeld and Haynes 2015), we then assessed whether distributed information across patterns of EEG spectral features could be used to decode different experiential states using MVPC. These analyses were further guided by our aim to evaluate the hypothesis that introspection confidence in experiential states reflects higher dissimilarity of the underlying neural representations.

## 2 Materials and Methods

### 2.1 Participants

Forty-six right-handed participants (28 females, age range: 18-43, *M*_Age_=25.9, *SD*=5.7; years of education [post-secondary school]: *M*_Yoe_=4.3, *SD*=2.2) with normal or corrected-to-normal vision provided written informed consent to volunteer in the study and were compensated £10 per hour. A sample size of 40 allowed us to detect paired-samples effects of *d*≥.45 (*α*=.05, 1-*β*=.80, two-tailed). We recruited 46 participants due to potential attrition and loss of participants because of insufficient numbers of trials for the different state responses. All participants self-reported proficiency in English (1=no proficiency, to 10=native speaker; *M*=9.2, *SD*=1.04). Seven participants were excluded due to technical issues during EEG data recording (*n*=1), or insufficient number of response types in the task (*n*=6; see section *2.5*), resulting in a final sample of 39 participants (25 females, age range: 18-43, *M*_Age_=25.6, *SD*=5.8, English language skills: [*M*=9.2%, *SD*=1.0]). The study was approved by the Research Ethics Committee of the Department of Psychology at Goldsmiths, University of London.

### 2.2 Materials

#### Audiobook listening task

This task consisted of participants listening to an audio version of Bill Bryson’s *A Short History of Nearly Everything* (2004), a general science book that has previously been used in mind wandering research (e.g. Smallwood, Nind, et al. 2009). Participants focused on a central white fixation cross on a grey background at a distance of approximately 90cm and listened (through speakers) to the audiobook in three 20min blocks (corresponding to chapters 7, 24, and 30 in counterbalanced order). During the task, participants were prompted via on-screen thought probes at pseudorandom intervals (30, 40, or 50s) to report on their experiential state: “Just before the probe, were you mind wandering?” (ES Judgement, response options: yes, no). If a participant responded in the affirmative, they were next prompted regarding state meta-awareness: “Just before the probe, were you zoning-out or tuning-out?” (MA Judgement, response options: tuning-out, zoning-out). Participants responded to both probes using a continuous visual analogue scale in which they made their binary judgement combined with an estimate of confidence in their response (ranging from completely not confident, to completely confident). Response options for both probes alternated sides randomly to control for response biases.

#### Audiobook listening assessment

A sequence of 20 true/false questions (corresponding to the content of the preceding block) were administered to participants after each block. Question order followed the presentation order of the information in the audiobook with each question corresponding to approximately 1 minute of content.

### 2.3 Procedure

After EEG preparation and general instructions, participants completed a battery of psychometric measures (to be reported elsewhere). Participants sat in a dimly lit room and first underwent a 5-min eyes-open resting state condition in which they focused on a central white fixation cross (1cm^2^) whilst their EEG was recorded and subsequently completed a self-report resting state measure (to be reported elsewhere).

Prior to completing the task, mind wandering was defined to participants as any thoughts that are not related to the material being presented (Christoff et al., 2009; Lindquist & McLean, 2011), and are usually internally focused. Participants were provided with examples of mind wandering, such as thoughts about past events, friends or significant others or concerns about an upcoming exam (Varao Sousa et al., 2013). Tuning-out was defined as a state in which one mind wanders and is aware whilst they are doing so whereas zoning-out was defined as a state in which one mind wanders and is unaware that they are doing so until they “catch” themselves.

The experimenter introduced the audiobook listening task and defined the response options in the ES and MA judgements and ensured that participants understood these options. Participants completed a 3min training block followed by three 20min experimental blocks (30 probes per block), resulting in 90 probes in total. After each block, participants completed the audiobook listening assessment. Blocks took approximately 25 mins to complete with the entire experiment lasting approximately 2.5 hours. The experiment was programmed and implemented in MATLAB® (2018a, The MathWorks, Inc., MA), using the Psychophysics Toolbox extensions (Kleiner et al., 2007).

### 2.4 Behavioural analyses

Participants’ data were segregated at the probe-level according to self-report in two ways for separate analyses: dichotomously (on-task vs. mind wandering) and trichotomously (on-task vs. tuning-out vs. zoning-out). Frequency (%) of each state report and performance on the audiobook listening assessment (accuracy [%]) were additionally computed at the block-level.

### 2.5 Electrophysiological data acquisition and analyses

EEG signals were recorded using a 64-Ag-AgCl electrode Biosemi ActiveTwo system. Electrodes were placed according to the International 10-20 system. Two electrodes placed on the participants’ earlobes were used as reference. Additional electrodes recorded right side vertical (VEOG) and bilateral horizontal (HEOG) electro-oculogram signals to be used for artefact detection and rejection. The recording was sampled at 512Hz for all participants.

Data pre-processing was implemented using the EEGlab toolbox in MATLAB (Delorme & Makeig, 2004). The average of the two earlobe electrodes was used as reference and the data were subsequently filtered with a high-pass filter at 0.5Hz and a notch-filter between 48-52Hz. We used the pop_eegfiltnew function in EEGlab, which applies a finite impulse response (FIR) filter to the data using an automatic filter order (3380 and 846, respectively). Bad electrodes (range: 0-2 across participants) detected during the recording, or via visual inspection of the raw data, were removed. Next, independent component analysis (ICA) was performed on the continuous data to detect eye-movement artefacts. IC scalp maps, spectra and raw activity were visually inspected to reject further artefacts (range: 1-5, *M*=3.15, *SD*=1.48) such as eye movements, further channel noise, and prominent muscle movements. Next, data from removed electrodes were replaced using spherical interpolation and all data were re-referenced using the average of the 64 channels.

For each participant, continuous data were next segmented into 14s epochs: −12 to 2s relative to probe onset. The time window of interest extended from −10 to 0s, but an additional 2s were included on each side to avoid edge artefacts in subsequent analyses; these data were omitted after time-frequency transformation. Further epoch exclusion was conducted via manual rejection based on visual inspection. Data were subsequently segregated into two conditions corresponding to the experiential states reported by the participants: on-task vs. mind wandering. Mind wandering states were further partitioned into two meta-awareness states: tuning-out vs. zoning-out. A 700ms epoch during the inter-stimulus interval after the probe response phase and start of the next trial was used as a baseline. A time-frequency transformation of the data was implemented by applying a Hanning window at 50ms steps to each 14s-long epoch and corresponding baseline segment for frequencies of 1 to 45Hz. Window length varied along the frequency dimension, with 7000ms at the lowest frequency (1Hz) decreasing linearly (time window=7/frequency) at each frequency bin, and 150ms for 45Hz. Trial-wise spectral power was averaged and then normalised by division of the baseline level (gain model; Grandchamp and Delorme 2011).

Participants varied in their mind wandering and state meta-awareness reports, resulting in different sample sizes and respective numbers of trials per state for each comparison. State-specific data that included fewer than 10% of probes (9 trials) were excluded from any analyses involving the respective state. Four main contrasts were implemented with variable trials (*M*±*SD*) and sample sizes [*N*]. After artefact removal and participant exclusions the number of trials per state for each of the four contrasts were: (i) on-task (39.7±15.0) vs. mind wandering (27.4±12.0) [*N*=39]; (ii) on-task (36.0±13.0) vs. tuning-out (20.5±9.8) [*N*=27]; (iii) on-task (35.5±13.7) vs. zoning-out (14.1±5.4) [*N*=25]; and (iv) tuning-out (22.0±13.4) vs. zoning-out (13.3±4.0) [*N*=21]. Insofar as the number of trials differed across states, we performed a series of control analyses in which significant contrasts were repeated with closer matching of the number of trials; these yielded similar results (see Control Analyses in **Supplementary Materials**).

### 2.6 Multivariate pattern classification analysis (MVPC)

#### 2.6.1 Time-varying MVPC

Complementing the univariate analysis of spectral power differences, we implemented a two-class MVPC for the four aforementioned two-state contrasts. Here, we hypothesized that information about the different experiential states would be shared across different frequency bands and electrodes. Insofar as this information could unfold over time, we used time-varying MVPC to investigate whether subjective reports about experiential states could be decoded from trial-wise EEG patterns of oscillatory activity across different frequency bands, and separately for different time points.

The features used for the decoding analyses were the trial-wise measures of spectral power for delta, theta, and alpha frequency bands for each of the 64 EEG channels. After pre-processing the data (see section *2.5*), a time-frequency transformation was implemented in the same way as in spectral analysis however data were neither baselined nor collapsed across trials. The spectral power values were then averaged separately for each frequency band (delta [2-3Hz], theta [4-7Hz], alpha [8-13Hz]). For the time-varying MVPC, these measures were additionally averaged in bins of 500ms within 10s epochs, resulting in 21 time bins. A linear support vector machine (SVM library for MATLAB; Lotte et al. 2007; Chang and Lin 2011) was trained to distinguish between classes (states) at each time bin. In order to examine whether single-frequency classification was superior or comparable to cross-frequency classification (multi-frequency model), we further performed MVPC analyses separately for each frequency band (reduced feature space). The multi-frequency model used data with 192 (64 electrodes x 3 frequency bands) dimensions for each trial, while the three single-frequency models (delta, theta, alpha) were comprised of 64-dimensional data per trial. Each of the implemented two-class MVPC analyses were balanced by matching the number of trials in each class using random trial selection. The matching of trials resulted in the following number of trials (*M*±*SD*) per class for each two-class decoding: (i) on-task vs. mind wandering (23.4±9.0, *N*=39); (ii) on-task vs. tuning-out (18.8±8.2, *N*=27); (iii) on-task vs. zoning-out (13.9±5.1, *N*=25); and (iv) tuning-out vs. zoning-out (12.6±3.6, *N*=21). MVPC was implemented using a 3-fold leave-one-out cross-validation procedure to assess the effectiveness of the model.

The individual empirical decoding accuracy was the average accuracy across the three folds.

Details on the participant- and group-level statistical analyses on time-varying MVPC decoding accuracies are described in section *2.7.3*.

#### 2.6.2 Time-averaged MVPC

An important consideration is that, although mind wandering processes could unfold over time, they are inherently not time-locked, and indeed our experimental design was not event-related. To account for the case that there would not be a reliable temporal representation of mind wandering states across trials, we further complemented the time-varying MVPC with a time-averaged MVPC approach. We collapsed the spectral power measures across time to assess whether we could still decode classes from the same features. An additional advantage of this second approach is that using time-averaged measures of spectral power in MVPC reduces the number of multiple comparisons. Spectral power at each time point was averaged across the whole epoch (−10 to 0s before probe onset). Similar to the time-varying analyses, the time-averaged MVPC was performed using the spectral power values for each frequency band and for the set of 64 channels as features. This was performed separately for the multi-frequency model (192 dimensions) and for each frequency band of interest (single-frequency models: 64 dimensions). This was carried out 11 times after partitioning the data into balanced classes 11 times, which aimed to obtain accuracy estimates more representative of the total trial sample. Classification parameters and trial amounts per class used for each two-class time-averaged MVPC were the same with time-varying MVPC (see section *2.6.1*). The individual empirical decoding accuracy was the average accuracy across the three folds and 11 analyses. Details on the participant- and group-level statistical analyses on time-averaged MVPC decoding accuracies are described in section *2.7.3*).

The general time-averaged MVPC was complemented with a method that allows assessing the activation patterns that describe the contribution of each channel or frequency band to the decoding of class-related information (Haufe et al., 2014). In the case of linear classifiers, we can obtain an estimate of the activation patterns **A** by projecting the extraction filters of the classifier, **W**, onto the channels as follows: **A ∝ Σ**_x_**W**, where **Σ**_x_ is the covariance matrix of the data. This calculation can be simplified by computing the covariance between the data (***x***) and hidden factors (***y***, vector of labels of each class): **A** ∝ **Σ**_x_**W**=*Cov*[***x****(n), **y**(n)*], where *n* is the number of trials (Haufe et al., 2014). The covariance *Cov*[***x****(n), **y**(n)*] is computed for each pattern dimension (i.e. each channel or frequency band) across the total number of training trials *n*, and separately for each fold (followed up by an average across folds). The amplitude and sign of the covariance matrix *Cov*[***x***(*n*), ***y***(*n*)] can be interpreted as the strength and polarity with which class-related information is reflected in each pattern dimension (channel or frequency band). Note, however, that MVPC is a multivariate analysis method, and thus inferences regarding univariate measures (e.g. individual channels) should be interpreted carefully. For multi-frequency models, we averaged the activation maps across frequency bands, to obtain a single topographic representation across 64 channels. The activation patterns associated with each frequency range were assessed using the single-frequency models.

Finally, we conducted correlation analyses between decoding accuracies derived from our time-averaged models and participants’ confidence in their respective reports. Correlation analyses were performed exclusively in a subset of the models, based on our EEG findings and previous research (see section *2.7.3*).

### 2.7 Statistical analyses

#### 2.7.1 Behavioural data

Confidence ratings between states were compared with paired-samples *t*-tests (two-tailed) and assessment performance was compared to 50% using a one-sample *t*-test. Associations between task-level mind wandering frequency and assessment accuracy were assessed by correlational analyses following automatic bivariate outlier (boxplot method) removal using the Robust Correlation toolbox in MATLAB (Pernet et al., 2013) in the computation of skipped correlations (Rousseeuw, 1984; Rousseeuw & Van Driessen, 1999; Verboven & Hubert, 2005). We report Spearman’s *r*_s_ for data that violated parametric test assumptions.

#### 2.7.2 Spectral analysis

Significant differences in the spectral power between different states were assessed by means of cluster-based paired permutation tests across participants (Maris & Oostenveld, 2007). Non-parametric cluster permutation tests were used separately for pre-specified oscillatory frequency bins (delta [2-3Hz], theta [4-7Hz], alpha [8-13Hz]). These analyses were undertaken in two phases. First, we calculated the observed test statistic for the respective contrast by: (i) conducting paired-samples *t*-tests comparing the two states at each data sample (frequency x channel x time); (ii) samples whose *t*-values were below threshold (α<.05) were selected and clustered in sets based on feature adjacency (spectral, spatial, and temporal); and (iii) *t*-values were summed to compute cluster-level statistics whose maximum served as the test statistic to evaluate state differences. The second phase entailed the same steps but this time the test statistic was computed for 500 permutations of randomly partitioned data in two subsets (Monte Carlo permutation test). These test statistics were compared to the observed test statistic (Maris & Oostenveld, 2007). The cluster-level significance value was set at two-tailed α<.025 with a minimum of 2 neighbouring channels constituting a cluster. This method controls for multiple comparisons by controlling the family-wise error rate at α=.05 We interpret effects in the range of .025<*p*<.030 as reflecting trends. For non-significant findings, we report *p*-values for the most prominent cluster. The analyses were conducted with the Fieldtrip toolbox in MATLAB (Maris & Oostenveld, 2007). Effect sizes were estimated using Hedges’s *g* and bootstrap 95% confidence intervals (CI, bias-corrected and accelerated method, 10000 samples [Efron 1987]), on power averages across frequency, time window, and electrode sites identified by cluster analyses, using the Measures of Effect Size Toolbox in MATLAB (Hentschke 2021).

#### 2.7.3 MVPC analyses

Statistical inference for the time-varying MVPC analyses was initially performed at the participant level, followed by group-level analysis. At the participant level, the null distribution for accuracy was generated by performing MVPC 500 times after randomly shuffling the class labels in the data. *P*-values were computed at each time bin as the proportions (%) of permutation accuracies that are greater than or equal to the observed decoding accuracy (mean accuracy of 3-fold cross-validation), yielding one *p*-value per time bin. We report the proportion of participants for which we observed significant decoding in at least one time bin. Next, at the group level, we aimed to identify the time bins showing statistically significant above-chance decoding accuracy, using a pairwise permutation test (Monte Carlo permutation test, 5000 iterations). Both at the participant- and group-levels, we corrected for multiple comparisons by controlling the false discovery rate (FDR) at .05 by using an adaptive two-stage linear step-up procedure (Benjamini et al., 2006). At the group-level, the corrected threshold *p*-value obtained from this procedure, *p*th, is given when multiple comparisons were performed.

Time-averaged MVPC analyses were performed in the same manner but were limited to a single averaged time bin and repeated 11 times. At the participant level, the null distribution for accuracy was computed by performing the analysis 500 times after randomly shuffling the class labels in the data. For each of the 11 repetitions, *p*-values were computed as the proportions (%) of permutation accuracies that are greater than or equal to the observed decoding accuracy (mean accuracy of 3-fold cross-validation), yielding one *p*-value per participant for each repetition. We report the range and mean percentage of participants across the 11 repetitions for which we observed significant (*p*<.05) decoding for each model. We then obtained the average empirical decoding accuracy per participant across the 11 analyses and used the results to conduct statistical analyses at the group level. Here we used as chance level the average accuracy of the permutation distributions (11 analyses) in each participant. Statistical assessment with a permutation test was performed by comparing these group mean accuracies and the estimated null distribution (Monte Carlo permutation test, 5000 permutations) to identify statistically significant decoding accuracy across participants at *p*<.05. Different models were compared using the Monte Carlo approach described above (paired permutation test).

Finally, we assessed associations between mean participant-level decoding accuracies (averaged across the 11 repetitions) and participants’ corresponding mean confidence ratings. Toward this end, confidence in ES and MA Judgements were averaged within participants for each state in order to obtain the average confidence for each class. These were then correlated at the group-level with decoding accuracies of the respective classification. To minimize the family-wise error rate, we selected a small number of models for this analysis based on the previous literature and our EEG data. For decoding of mind wandering vs. on-task states and tuning-out vs. on-task states, we examined the associations between decoding accuracies in the multi-frequency model and confidence ratings. For zoning-out vs. on-task states and zoning-out vs. tuning-out decoding we examined correlations between confidence ratings and the delta and theta models’ decoding accuracies. As described above, bivariate outliers were removed in the computation of skipped Pearson correlations (Rousseeuw, 1984; Rousseeuw & Van Driessen, 1999; Verboven & Hubert, 2005). For discrepancies between 95% confidence intervals (95% CIs) and *p*-values, correlations are interpreted as non-significant.

## 3 Results

### 3.1 Characteristics of mind wandering, state meta-awareness and introspection confidence

During the audiobook listening task, participants (*N*=39) reported mind wandering (*M*%±*SD*) on 39.7±17.9 of the probes, with stable rates across blocks (block 1: 41.2±20.6; block 2: 40.9±21.6; block 3: 36.9±19.0). Among mind wandering states, participants reported tuning-out (56.6±19.0) more often than zoning-out (43.4±19.0). Participants varied (range, *M*%±*SD*) in their confidence for ES judgements (14.8-91.4, 61.7±7.7), and displayed less confidence in mind wandering reports (11.3-90.0, 53.2±19.9), than in on-task reports (8.6-94.2, 64.4±19.0), *t*(38)=4.78, *p*<.001, *g*=.57, [0.32 0.90]. Specifically, on-task reports were rated with significantly higher confidence than both tune-out (*t*(38)=2.39, *p*=.02, *g*=.34, [0.08 0.67]) and zone-out reports (*t*(38)=3.80, *p*<.001, *g*=.58, [0.28 0.96]). Participants were moderately confident in their MA judgments during mind wandering states (16.7-92.0, 58.1±6.3), with numerically, albeit non-significantly, greater confidence in tune-out (17.2-92.3, 58.0±18.8) than in zone-out (6.9-90.5, 52.3±21.8) reports, *t*(38)=1.55, *p*=.13, *g*=.27, [−0.06 0.64]. These rates are similar to previous research (Christoff et al. 2009; Seli et al. 2015; Varao Sousa et al. 2013) and demonstrate variability in experiential states, state meta-awareness, and introspection confidence during the task.

### 3.2 Audiobook listening assessment and mind wandering frequency

Accuracy on the assessment averaged across blocks (*M%*±*SD*: 74.5±10.0) was above chance performance (50%, one sample *t*-test: *t*(38)=15.3, *p*<.001, *g*=2.44, [1.84, 3.59]). Performance was comparable across blocks (block 1: 72.3±12.0, block 2: 74.6±14.4, block 3: 76.5±12.1), suggesting stable motivation throughout the task. Mind wandering frequency reliably significantly correlated negatively with assessment accuracy in the last two blocks (block 1: *r*=-.29 [95% CI: −.53, .02], *p*=.079 (*p*=.017, with outliers), block 2: *r*_s_=-.43 [−.68, −.10], *p*=.009 (*p*<.001, with outliers), block 3: *r*=-.41 [−.68, −.11], *p*=.011 (*p*=.011, with outliers), thereby providing an indirect behavioural validation of participants’ self-reports and corroborating previous research (Schooler et al. 2004; Boudewyn and Carter 2018).

### 3.3 Oscillatory characteristics of mind wandering and state meta-awareness

As expected, the cluster-based permutation test revealed greater alpha power during mind wandering than on-task states (**Figure 1**). The analysis revealed two temporally-adjacent clusters just prior to probe onset, *p*=.004, *g*=0.56 [0.32, 0.92]; *p*=.024, *g*=0.50 [0.29, 0.80]. Both effects were topographically diffuse and most pronounced over bilateral frontocentral and right posterior sites. Similarly, we observed greater alpha power during tuning-out than on-task states in a single cluster, *p*=.008, *g*=0.66 [0.37, 1.05] (**Supplementary Figure 1**). This effect was also close to probe onset and was primarily observed over fronto-central and parieto-occipital regions. There were no other significant differences between states in the other frequency bands (**Supplementary Figure 2**): for on-task vs. mind wandering, the analysis did not yield any significant clusters for delta, *p*=.24, or theta power, *p*=.40. Similarly, delta and theta power did not significantly differ between tuning-out and on-task states (**Supplementary Figure 2**, delta: *p*=.23; theta: *p*=.20).

**Figure 1.**
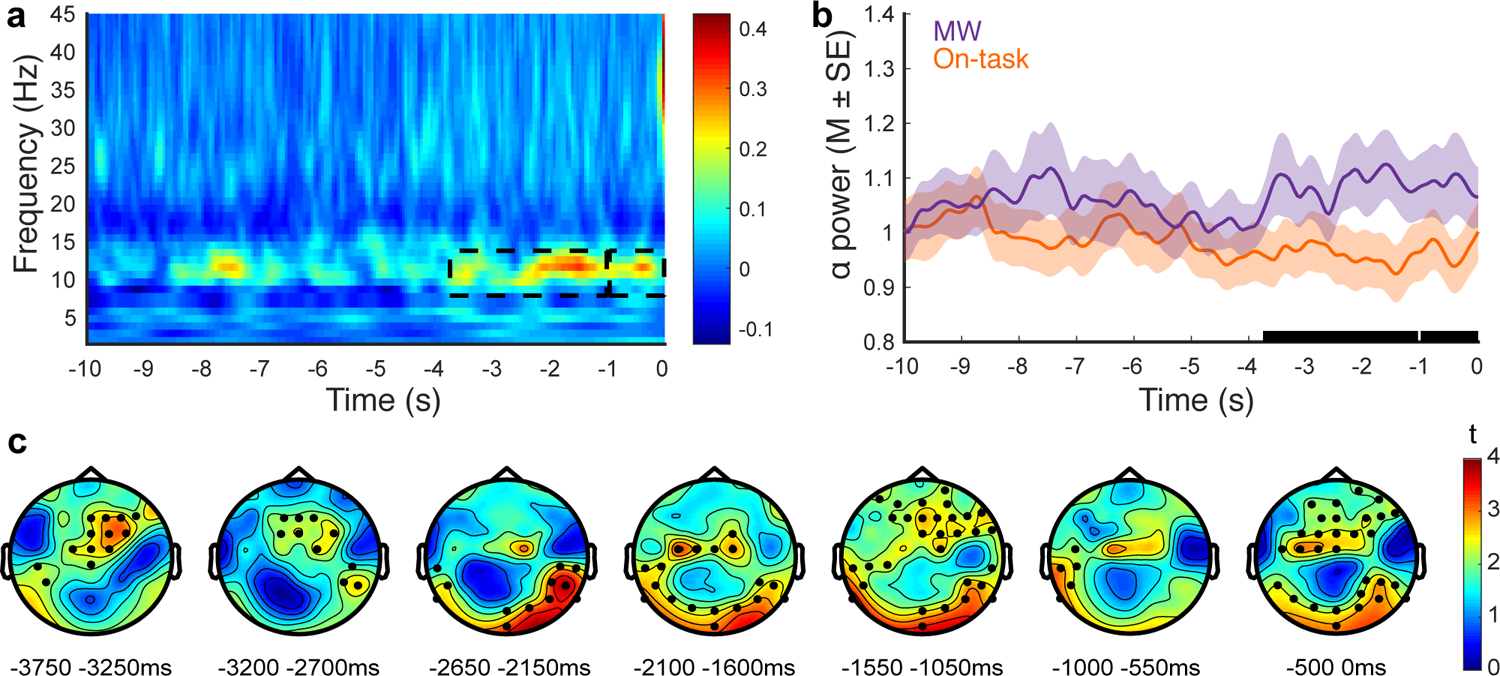
Oscillatory differences between states (MW – on-task, *N*=39) as a function of time relative to probe onset (0s). a) Time-frequency decomposition averaged across electrode sites. Broken black rectangles denote spectrotemporal clusters reflecting significant state differences (*p*<.025, two-sided cluster-based permutation test). b) Alpha (8-13Hz) spectral power averaged over the electrode sites of the two clusters (significance denoted by black bars on the x-axis). c) Topography of the clusters at different 500/550ms sub-windows (black markers denote electrodes that were present on at least 50% of samples in each time window). MW = Mind Wandering.

In line with the foregoing results, zoning-out (unaware mind wandering) states were characterized by greater power than on-task states in delta, theta, and alpha bands in two distinct time windows (**Figure 2**). Alpha power was greater for zoning-out than on-task states in a short interval early in the epoch, *p*=.004, *g*=0.75 [0.51, 1.06], and a long interval just prior to probe onset, *p*=.002, *g*=0.86 [0.57, 1.28]. Both effects were larger in magnitude than the comparisons between on-task and mind wandering and tuning-out states and topographically diffuse but strongest over bilateral frontal and posterior sites. Similarly, theta power was greater in zoning-out than on-task states in a single window overlapping with the late alpha effect, *p*=.010, *g*=0.83 [0.54, 1.24]; this effect was larger in right frontal electrodes but over time shifted to left temporo-parietal sites. A second theta band cluster overlapped in time with the early alpha cluster but did not achieve significance despite a large effect size, *p*=.028, *g*=0.76 [0.50, 1.14]. Zoning-out states were also associated with greater delta power than on-task states in two clusters that were temporally coincident with the foregoing effects but with substantially larger effect sizes, *p*=.018, *g*=1.43 [1.04, 2.16]; *p*=.018, *g*=1.16 [0.74 1.73]. These effects were topographically more focal and largely restricted to midline central electrodes.

**Figure 2.**
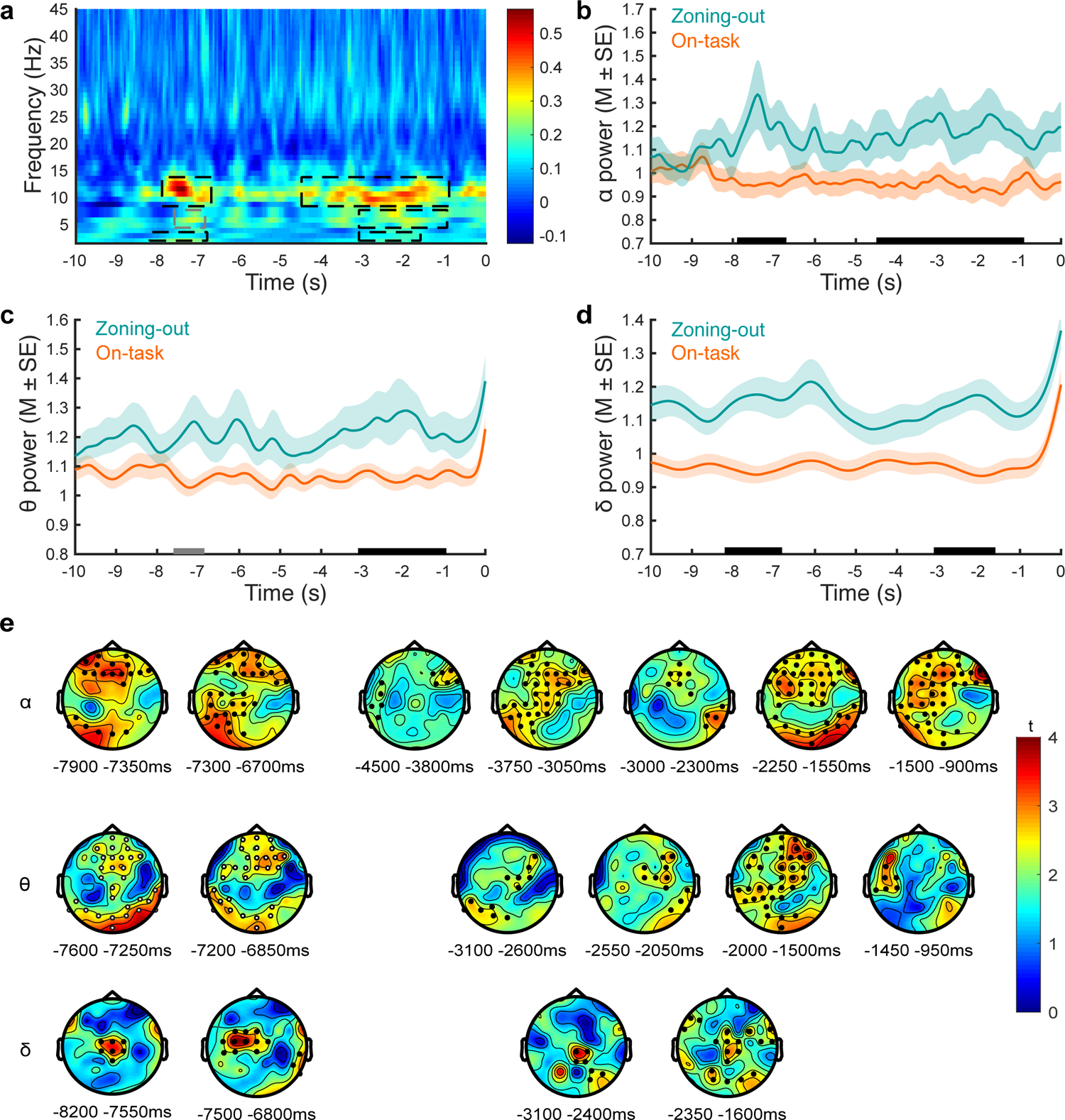
Oscillatory differences between states (zoning-out – on-task, *N*=25) as a function of time relative to probe onset (0s). a) Time-frequency decomposition averaged across electrode sites. Broken black rectangles indicate spectrotemporal clusters reflecting a significant difference (*p*<.025, two-sided cluster-based permutation test) and the grey rectangle indicates a trend-level (.025<*p*<.05) cluster. b, c, d) Alpha (8-13Hz), theta (4-7Hz) and delta (2-3Hz) spectral power averaged over the electrode sites of the clusters (black bars=significant, grey bar=trend). e) Topography of the clusters at different sub-windows within the cluster (black markers denote electrodes that were present on at least 50% of samples in each time window, white electrodes mark topography of the trend-level effect).

Zoning-out states were also characterized by greater theta power than tuning-out states in a single cluster close to probe onset at a trend-level of significance, *p*=.026, *g*=0.86 [0.45 1.41] (**Supplementary Figure 3**). This effect was most pronounced in parieto-central electrodes. Failure to reach significance is plausibly due to the reduced sample for this analysis (*N*=21) due to MW trial partitioning. There were no other significant effects in the other frequency bands (**Supplementary Figure 2**). Specifically, tuning-out and zoning-out states did not differ in delta (*p*=.05) or alpha power (*p*=.16). Similarly, exploratory analyses of beta power did not yield significant differences between any ES or MA states (see **Supplementary Figure 4**).

### 3.4 Multivariate pattern classification analyses

The univariate analyses of the EEG signals managed to identify spectral and spatio-temporal differences between states. This analysis, however, failed to demonstrate a robust and significant difference between aware and unaware mind wandering. In the following, we present the results for two sets of analyses: the time-*varying* MVPC analyses performed in the epoch before probe onset (−10 to 0s; 21 bins of 500ms) and the time-*averaged* MVPC analyses performed after collapsing temporal information for decoding across the entire epoch.

#### 3.4.1 Time-varying MVPC

For each two-class (state) comparison, we tested four models that used as features the spectral power in the 64 channels in the delta, theta and alpha frequency bands (multi-frequency model) and in these three bands separately (single-frequency band models). At the participant-level, we report the percentage of participants for which significant decoding was obtained in at least one bin.

The multi-frequency model (trials per class: 23.4±9.0) significantly decoded mind wandering from on-task states in 26% of participants, with similar proportions in single-frequency models: delta: 23%; theta: 15%; and alpha: 26% (**Supplementary Figure 5a)**. At the group level, both the multi-frequency and the single-frequency models were able to decode mind wandering and on-task states with significant accuracies observed at various bins in the window (**Figure 3a**). Similarly, the multi-frequency model (trials per class: 18.8±8.2) significantly decoded on-task from tuning-out states in 22% of participants although the single-frequency models displayed superior decoding: delta: 44%; theta: 30%; alpha: 33% (**Supplementary Figure 5b)**. Group-level classification was significant in four bins in the multi-frequency model with similar results for theta and alpha, but not delta (**Figure 3b**). Significant decoding accuracy between zoning-out and on-task states (trials per class: 13.9±5.1) was observed in 24% of participants for the multi-frequency model, with comparable or superior accuracy in the single-frequency models (delta: 44%; theta: 36%; alpha: 24%; **Supplementary Figure 5c**). Group-level significant decoding was found for the multi-frequency model and the single-frequency models (**Figure 3c**). The theta and delta effects were spread throughout the window with the delta model exhibiting significant decoding accuracy across the full window. Tuning-out and zoning-out states (trials per class: 12.6±3.6) were decoded in 14% of participants in the multi-frequency model, with superior decoding in the single-frequency models (delta: 24%; theta: 19%; alpha: 29%, **Supplementary Figure 5d**). At the group-level, the multi-frequency and single-frequency models showed significant decoding although this was restricted to one bin in the former and generalized throughout the window in the latter (**Figure 3d**). These results indicate that decoding for all two-class comparisons was significant in several time bins for models using both the multi-frequency and the reduced feature space (single-frequency) suggesting that the individual frequency bands alone are sufficient to decode experiential states and state meta-awareness.

**Figure 3.**
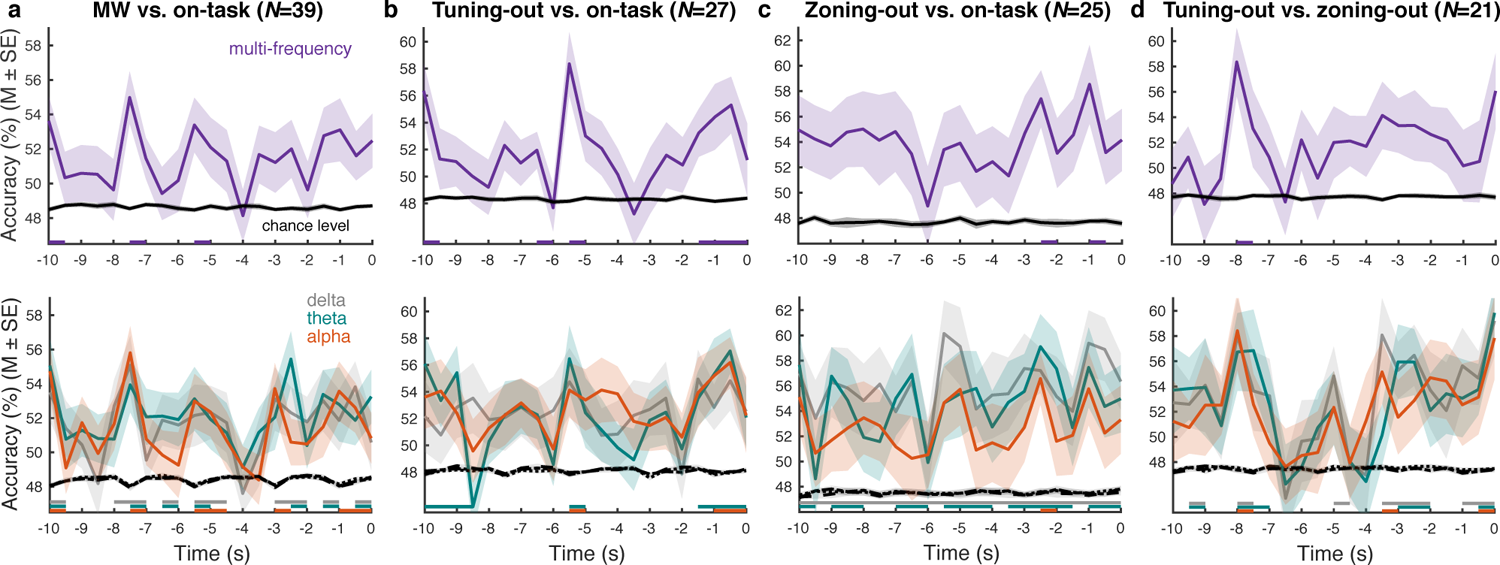
Time-varying MVPC (Multivariate Pattern Classification analysis): Group decoding accuracy across participants in multi-frequency (**top panels**) and single-frequency (**bottom panels**) models for four different state contrasts. The null distribution was obtained by 5000 random permutations after shuffling the labels (black lines). Above chance decoding (relative to the permutation distribution) is denoted by horizontal x-axis coloured markers (based on an FDR correction; see **Supplementary Table 1**). MW = Mind Wandering.

#### 3.4.2 Time-averaged MVPC

At the participant-level, the multi-frequency model significantly decoded mind wandering from on-task states in (*M*±*SD*) 19±6% (range:10-28%) of participants (**Figure 4a**). Individual-frequency models decoded states in similar proportions of participants: delta: 13±3% (10-21%); theta: 15±5% (8-26%); alpha: 22±4% (15-26%), and did not significantly differ, *p*>*pth* (**Figure 5e**). At the group-level (**Figure 5a**), the multi-frequency model displayed significant decoding accuracy (M%±SE: 54.72±1.47, *p*=0), as did single-frequency models: delta (52.85±1.11, *p*<.001), theta (53.23±1.18, *p*<.001), and alpha (55.11±1.31, *p*=0). The multi-frequency model decoded tuning-out from on-task states in 14±7% (4-30%) of participants (delta: 12±4% [4-19%]; theta: 14±6% [11-30%]; alpha: 12±6% [4-22%]), with no significant differences between models, *p*>*pth* (**Figure 5f**). At the group-level (**Figure 5b**), the multi-frequency model displayed significant decoding accuracy (53.93±1.51, *p<.*001), as did the single-frequency models (delta: 52.52±1.30, *p*<.001, theta: 54.77±1.26, *p*=0, alpha: 52.61±1.30, *p*<.001).

**Figure 4.**
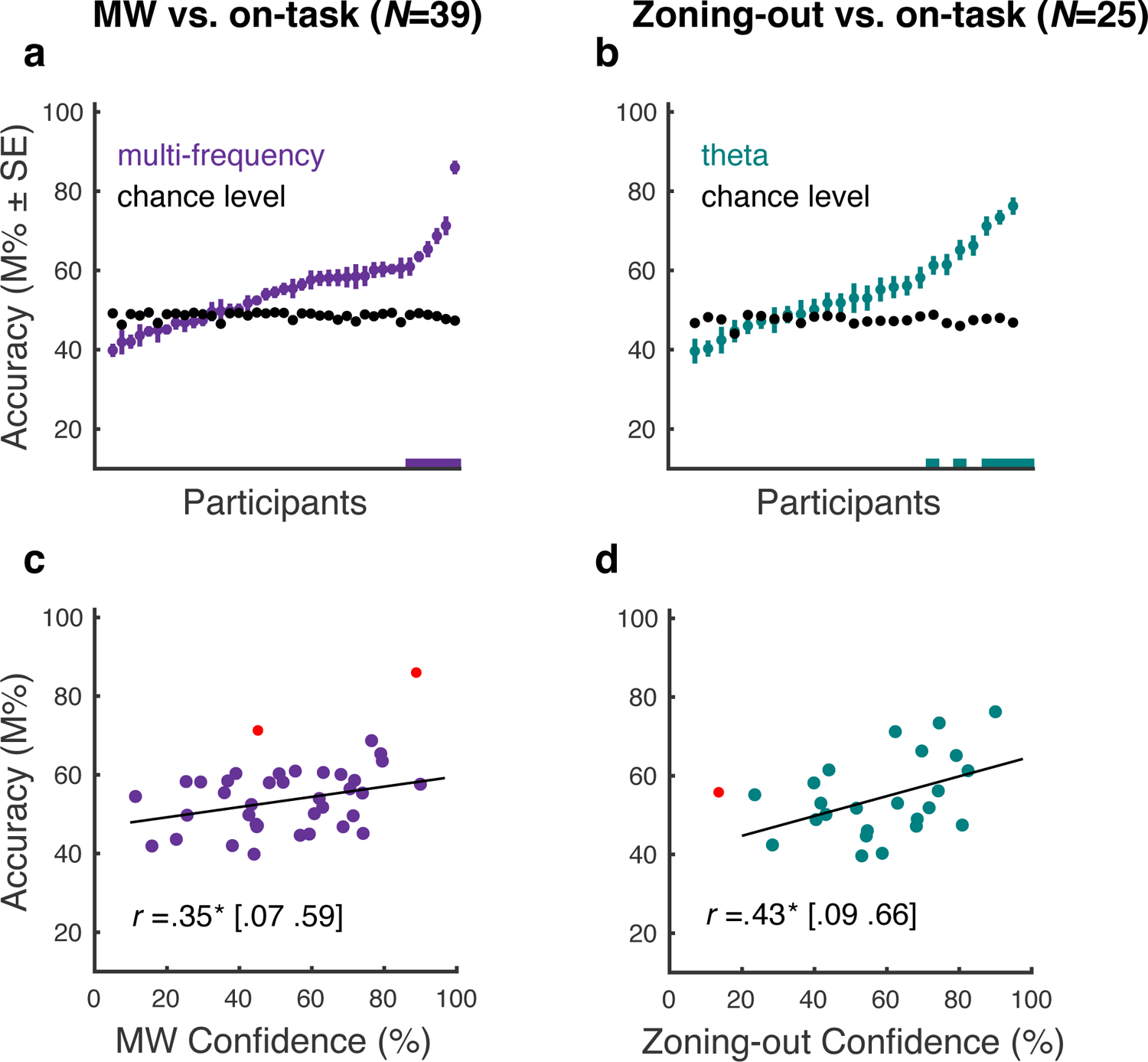
Time-averaged MVPC (Multivariate Pattern Classification analysis): (**a-b**) Decoding accuracy in individual participants relative to permutation distributions: chance level obtained by 500 random permutations after shuffling the class labels (upper panel). Participants for which above chance decoding was observed (relative to the permutation distribution) are denoted by horizontal x-axis coloured markers. (**c-d**) Scatterplots of participant-level mean decoding accuracies and mean judgment confidence ratings. Note that the empirical decoding accuracies in a few participants were below their respective chance level, which is often the case in the type of MVPC analysis we conducted (using the same two classes for training and testing data, Allefeld et al., 2016). What cannot be below chance level is the true single-subject accuracy, as it measures the amount of information (Allefeld et al., 2016). Red markers denote bivariate outliers; square brackets denote Bootstrap 95% CIs. MW = Mind Wandering. * *p*<.05

**Figure 5.**
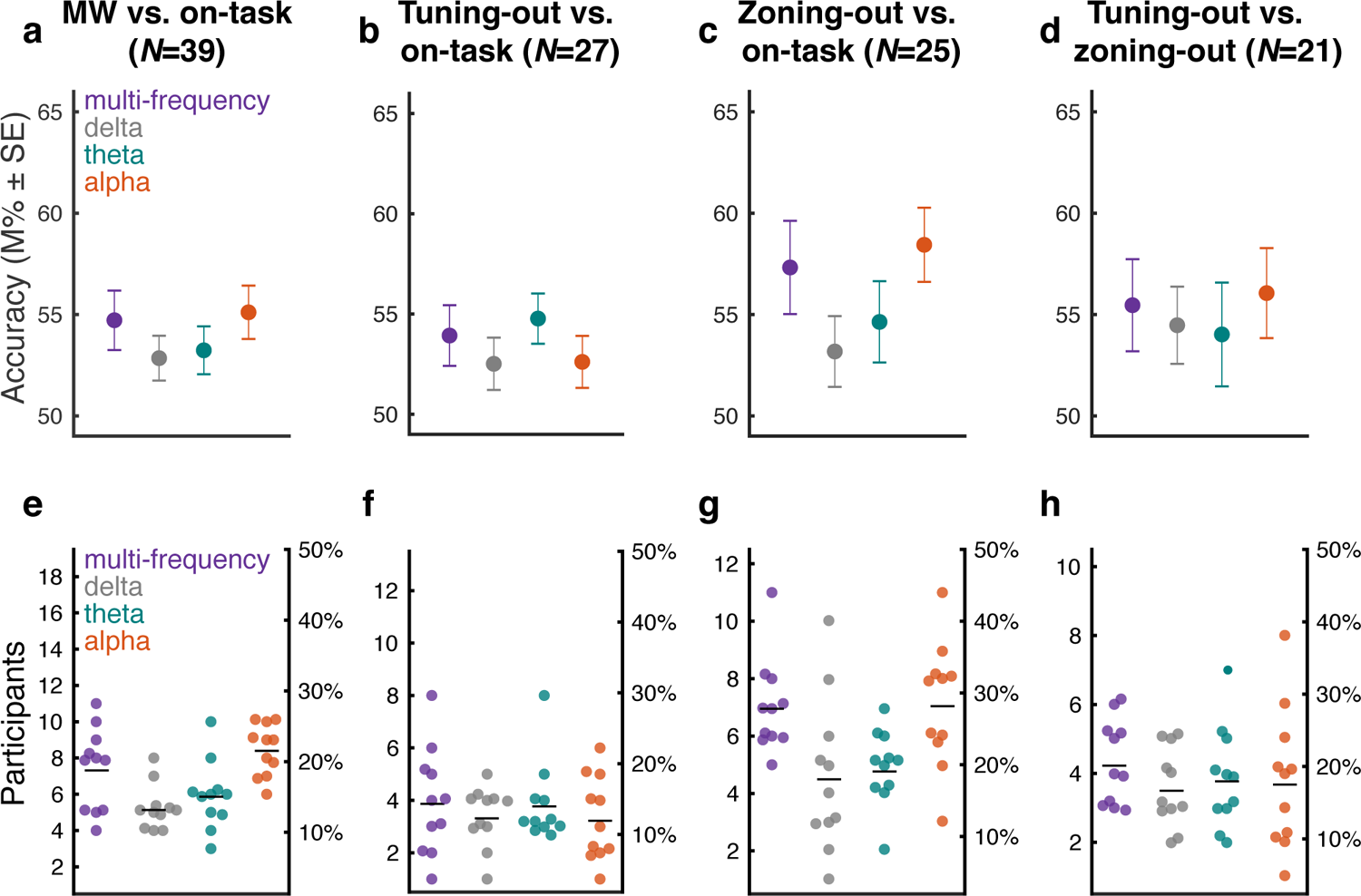
Time-averaged MVPC (Multivariate Pattern Classification analysis): **(a-d)** Group decoding accuracy across participants in multi-frequency and single-frequency models for four different state contrasts. (**e-f**) Participants (count and percentage) with significant (*p*<.05) decoding for each of the 11 MVPC repetitions. Black lines indicate the mean across the 11 repetitions. MW = Mind wandering.

Zoning-out from on-task states were significantly decoded in 28±6% (20-44%) of participants. Performance varied across single-frequency models (delta: 18±11% [4-40%]; theta: 19±5% [8-28%, **Figure 4b**]; alpha: 28±9% [12-44%]). Both the multi-frequency and alpha models were significantly better than the delta, *p*=.004, *p*=.003 and theta models, *p*=.049, *p*=.035 (**Figure 5g**). At the group-level (**Figure 5c**), the alpha model showed the greatest decoding accuracy (58.44±1.83, *p*=0), albeit with comparable accuracy in the multi-frequency model (57.32±2.30, *p*=0). The delta (53.18±1.75, *p*=.002) and theta (54.64±2.01, *p*=.001) models were also significant.

Meta-awareness of mind wandering (zoning-out [unaware] vs. tuning-out [aware]) was significantly decoded in 20±6% (14-29%) of participants by the multi-frequency model. Single-frequency models decoded states in similar proportions of participants (delta: 17±5% [10-24%]; theta: 18±7% [10-33%]; alpha: 18±10% [5-38%]), and these differences were not significant, *p*>*pth* (**Figure 5h**). At the group-level (**Figure 5d**), all models showed comparable decoding accuracies (multi-frequency [55.46±2.27, *p*=.002], delta [54.47±1.91, *p*<.001], theta [54.02±2.56, *p*=.016], alpha [56.06±2.22, *p*=0]). Owing to the fluctuating nature of mind wandering states, we expected that time-varying MVPC would reveal less consistent individual and group-level effects than time-averaged MVPC. However, the results did not suggest any substantial differences between the two approaches.

To further uncover the activation patterns that represent the expression of class-decoding information in each channel, we visualised the topography of the activation patterns reconstructed from the classification filters, following Haufe and colleagues (2014). The polarity of the values in the activation map denotes the effect *direction* of the class-related information represented in each channel. For decoding on-task and mind wandering states (**Figure 6**), this analysis revealed that, for the theta model, class-related information was strongly present at frontal electrodes across participants and was associated with positive activation values. In the delta model, posterior electrodes represented class-related information more strongly, also in association with positive values. By contrast, pronounced negative values were observed for the multi-frequency and alpha models at left parietal and fronto-central regions, suggesting that these electrode regions represented class-related information more strongly (with a negative direction of the effect). The topography of the reconstructed activation maps for the additional MVPC analyses was partially different from the patterns obtained for on-task vs. mind wandering decoding (see supplementary results in **Supplementary Materials**, and **Supplementary Figure 6**).

**Figure 6.**
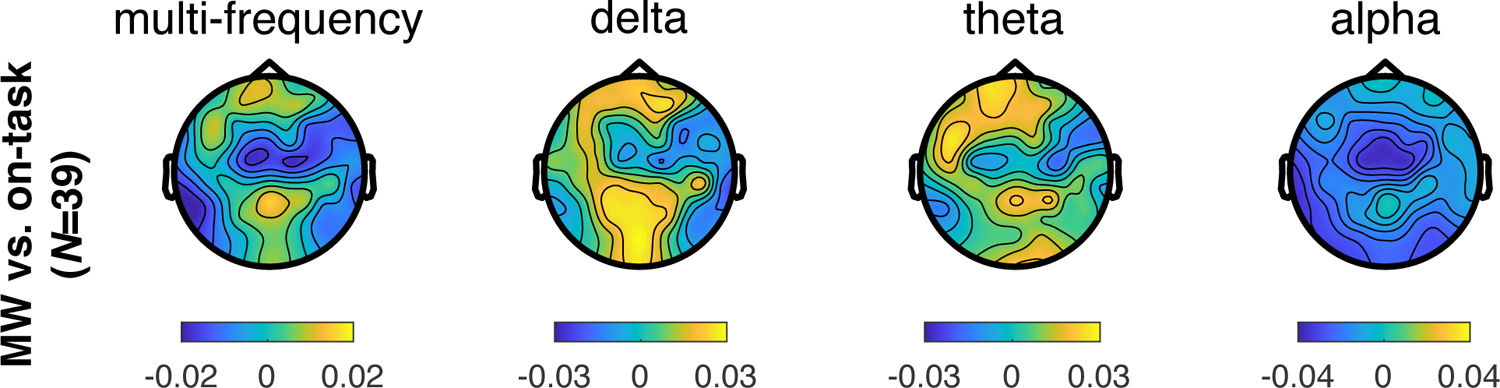
Activation patterns indicative of the strength and polarity with which class-related information is reflected in each pattern dimension (electrode site) for the time-averaged MVPC (Multivariate Pattern Classification analysis). Each column displays activation patterns for each model (multi-frequency, delta, theta, alpha) and for mind wandering vs. on-task classification. Activation values have arbitrary units.

#### 3.4.3. Association between decoding accuracy and introspection confidence

Our final set of analyses were motivated by the hypothesis that introspection confidence reflects conscious accessibility to experiential states. Toward this end, we evaluated the predictions that individual decoding accuracies in select models would positively correlate with confidence in experiential state and state meta-awareness judgments. We computed correlations between decoding accuracies of the multi-frequency model for on-task vs. MW and on-task vs. tuning-out states, and decoding accuracies of theta and delta models for zoning-out vs. on-task and zoning-out vs. tuning-out, with judgement confidence in the respective state.

In support of our overarching prediction, decoding accuracies for MW vs. on-task (multi-frequency model) positively correlated with ES judgement confidence ratings of mind wandering, *p*=.034 [0.07, 0.59] (**Figure 4c**, [*p*=.011, with outliers]), but not with on-task ES judgement confidence, *r*=.23, *p*=.18 [95% CI: −0.09 0.52] (*p*=.49 with outliers). Accuracies for the multi-frequency model decoding between on-task and tuning-out states did not correlate significantly with ES judgement confidence (on-task): *r*=.004, *p*=.99 [−0.34, 0.38] (*p*=.81 with outliers) or MA confidence (tuning-out): *r*=-.02, *p*=.92 [−0.48, 0.34]. Accuracies for the theta model decoding between on-task and zoning-out states did not correlate significantly with ES judgement confidence (on-task) *r*=.01, *p*=.95 [−0.30, 0.30] (*p*=.48 with outliers), but did correlate with MA confidence (zoning-out), *p*=.034 (**Figure 4d**, [*p*=.067 with outliers]). We also found that delta model decoding accuracies between on-task and zoning-out states did not correlate with ES confidence (on-task) *r*=-.09, *p*=.67 [−0.63, 0.04] (*p*=.39 with outliers) and MA confidence (zoning-out), *r*=.32 *p*=.12 [0.04, 0.57]. Accuracies for the theta model decoding state meta-awareness did not correlate significantly with MA confidence for tuning-out states, *r=*.08, *p*=.73 [−0.49, 0.44], nor with MA confidence for zoning-out states, *r*=.02, *p*=.95 [−0.40, 0.40]. Delta model decoding accuracies between tuning-out and zoning-out states did not correlate with MA confidence (tuning-out: *r*=.10, *p*=.66 [−0.27, 0.40], zoning-out, *r*=.35 *p*=.12 [−0.13, 0.66]).

## 4 Discussion

Using EEG and an ecological listening task, this study investigated the neural oscillatory dynamics of mind wandering and meta-awareness states. Mind wandering was reliably characterized by greater alpha power than on-task states with more prominent effects in this band and both delta and theta bands for unaware mind wandering. Consistent with the notion that mind wandering is more pronounced when one lacks meta-awareness (Christoff et al. 2009), moment-to-moment variations between unaware mind wandering and on-task states were the most reliably decoded via multivariate pattern classification. Critically, we found that decoding accuracy in the classification of different experiential states predicted confidence in the corresponding state judgments. This is consistent with the proposal that confidence indexes metacognitive access to experiential states.

### Mind wandering as external inattention and internal focus

As in previous research demonstrating the impact of mind wandering on comprehension (e.g. Boudewyn and Carter 2018), self-reported mind wandering was associated with poorer recall in the listening task. One of the principal results of this study was that power in the alpha frequency band was greater during mind wandering than on-task states, replicating previous work (Macdonald et al. 2011; Baldwin et al. 2017; Boudewyn and Carter 2018; Compton et al. 2019). This effect was specific to the alpha band and generalized across meta-awareness states suggesting that elevated alpha power is a frequency-specific but generalized neurophysiological characteristic of mind wandering. Alpha power differences had a relatively diffuse distribution and were concentrated in frontal and posterior sites. The broad distribution of alpha power is consistent with a previous study on mind wandering during a listening task and commensurate with alpha activity during auditory language comprehension (Boudewyn & Carter 2018, see also Compton et al., 2019). Posterior alpha has been extensively linked to mind wandering (Macdonald et al. 2011; Baldwin et al. 2017; Boudewyn and Carter 2018; Compton et al. 2019), as well as internally-oriented states more generally (Hanslmayr et al., 2011). Our findings are in line with studies showing elevated alpha power when attention is focused internally (Cooper et al. 2003) as well as multiple lines of evidence suggesting that mind wandering is associated with decay of perceptual processing (Smallwood et al. 2008; Barron et al. 2011). These results suggest that greater alpha power reflects detachment from the external world and a shift towards internal processing (Smallwood and Schooler 2015).

Our findings suggest that unaware mind wandering is more divergent from on-task states than episodes of mind wandering with awareness (Christoff et al., 2009). Although both were characterized by higher alpha power compared to on-task states (Boudewyn and Carter, 2018), unaware mind wandering was additionally associated with greater delta and theta power, particularly in right frontal and parieto-central sites, respectively. Aware and unaware mind wandering had suggestively distinct oscillatory features thereby implying that state meta-awareness represents a dimension of attention that is orthogonal to the direction of attention. These findings may help to explain previous reports of elevated delta and theta during mind wandering in self-caught paradigms (Braboszcz & Delorme, 2011; van Son et al., 2019), which in our view almost exclusively index unaware mind wandering, since they investigate EEG correlates seconds before participants realize they are mind wandering.

The observation of elevated delta and theta power during unaware mind wandering aligns with previous research on the cognitive correlates of oscillatory activity in these bands. Theta oscillations are suggested to be involved in cognitive control (Cavanagh & Frank, 2014), working memory (Klimesch, 1999; Mitchell et al., 2008) and conflict detection (Cohen, 2014), processes that could be implicated in appropriate selection between multiple simultaneous thoughts related to the processing of current concerns during unaware mind wandering. This aligns with hypothesised parallels between mind wandering and meditative states related to moment-to-moment navigation through mental objects (Vago & Zeidan, 2016). In particular, our finding of higher theta power during unaware than aware mind wandering states is potentially congruent with higher frontal midline and temporo-parietal theta during meditative states characterised by deeper absorption (DeLosAngeles et al., 2016) and thus suggest potential links between absorption and zone-outs.

Delta frequency contributions have been revealed during increased focus on internal processing and pertinent inhibition of interference (Harmony, 2013). Delta and theta activity could thus reflect the involvement of memory in self-related processing during self-generated thoughts in the context of unaware mind wandering episodes. The complementary role of delta activity might be to preserve internal processing during unaware mind wandering by inhibiting external interference (Harmony, 2013; Harmony et al., 1996). Accordingly, our findings may align with the proposal that mind wandering recruits processes to ensure that one’s internal train of thought is maintained (Smallwood et al., 2012; Smallwood, Fishman, et al., 2007).

### Decoding of experiential states

Although our data were not event-related, our time-varying multivariate classifier was able to trace mind wandering in several temporal segments within the 10 second time window preceding experiential state probes. Moreover, MVPC allowed us to decode experiential states from oscillatory activity at both participant- and group-levels, highlighting the utility of spectral measures coupled with machine learning in decoding mind wandering (Groot et al. 2021; Jin et al. 2019) and state meta-awareness. The time-varying and time-averaged analyses did not reveal substantially different results. In both analyses, all models decoded mind wandering from on-task states, including in approximately one-quarter of participants (multi-frequency, alpha) and with comparable classification accuracies. Similarly, aware mind wandering was decoded from on-task states in both analyses, with models using power in the delta and alpha bands for classification showing the weakest performance. It is potentially notable that the time-varying analyses yielded significant decoding between unaware mind wandering and on-task states in multiple time windows, particularly for the delta and theta models, which decoded the two experiential states across almost the entire epoch. By contrast, the time-averaged MVPC delta and theta models showed a weaker classification performance compared to the multi-frequency and alpha models which achieved the highest decoding accuracy at the group- and participant-levels. Although, insofar as all models achieved significant decoding, our analyses do not suggest that decoding between these two states is frequency specific per se. Finally, the time-averaged multi-frequency MVPC decoded aware from unaware mind wandering states in 20% of participants on average with comparable decoding accuracy values across all models.

Taken together, these results suggest that information pertaining to state meta-awareness might be distributed across different frequency bands including slow oscillations – as shown in the higher prevalence of significant decoding in the sample. However, our analyses did not reveal any robust evidence for frequency specificity and thus it seems that there is not a specific oscillatory pattern that contains more information about mind wandering and state meta-awareness. Assessment of the topography of the decoding results (Haufe et al., 2014) highlighted that the single-frequency models comprise differential activation patterns representing class-related information regarding experiential and meta-awareness states. Insofar as these analyses are purely descriptive, we should exercise caution in interpreting the topographic results, however these analyses suggested that class-related information tended to be strongly represented in fronto-central and posterior regions which broadly aligns with the effects observed in the cluster-based analysis.

We further corroborate that unaware mind wandering is more dissimilar at the neural level to external attention (on-task states) (Christoff et al. 2009). Combined with our MVPC results decoding aware from unaware mind wandering, we confirm that state meta-awareness should be considered as an important dimension of mind wandering in future studies, with evidence for disparate neural substrates for aware and unaware mind wandering. Collectively, MVPC trained on EEG-extracted features reliably decoded different experiential states both at participant and group levels. Discrepancies between group-level and participant-level classification is in line with research showing the utility of using individualized markers in decoding mind wandering (Dhindsa et al., 2019). Even though our task involved passive listening, one possibility is that residual eye movements could have contributed to the decoding. Eye movements can have confounding effects in neural decoding, particularly within visual tasks that involve active viewing (Thielen et al., 2019).

### Introspection confidence

Previous research investigating confidence suggests that self-reports of mind wandering characterised by higher confidence constitute more accurate evaluations of one’s experiential states (Seli et al. 2015). Our results build upon this, showing that confidence in experiential state judgments tended to map onto state meta-awareness: participants reported the greatest certainty for on-task episodes, were less confident in aware mind wandering, and the least for mind wandering episodes without awareness, although confidence ratings between the latter two were not significantly different. Future work could investigate how confidence relates to other prominent dimensions of mind wandering, such as intentionality (Seli et al., 2016, 2017). Participant-level decoding of experiential states allowed us to evaluate the prediction that introspection confidence would be positively associated with decoding accuracy. Indeed, confidence in different experiential state and state meta-awareness judgments reliably correlated with individual differences in MVPC decoding accuracies. In particular, confidence correlated with cross-frequency decoding accuracy in classifying mind wandering from on-task states. In addition, consistent with findings implicating theta activity in metacognition (Wokke et al., 2017), we also found that confidence was associated with decoding of unaware mind wandering from on-task states accuracy in the theta model.

Collectively, these results demonstrate that confidence in one’s experiential states is positively related to the multivariate decodability of these states with implications for the neural bases of experiential state confidence. Our results align with previous findings (Seli et al., 2015) suggesting that high confidence levels reflect a more accurate assessment of one’s experiential state reflected in stronger coupling between mind wandering reports and well-established impacts on behaviour. We extend this notion and provide evidence that higher confidence may reflect greater dissociation of experiential states at the neurophysiological level. In MVPC, higher decoding accuracy denotes better discriminability or separation between EEG patterns associated with each experiential state class. Although this discriminability refers only to pattern analysis, it is possible that more dissociable or distinct neural patterns are metacognitively represented and therefore reflected in individuals’ confidence ratings. One interpretation of these findings thus is that states accompanied by high confidence are quantitatively more intense or salient and thus characterized by superior conscious access that is grounded in or related to underlying neurophysiological discriminability. Alternatively, this relationship might be attributable to participants with low confidence displaying weak experiential state discriminability, i.e., they have relatively poor metacognitive access in their experiential states, associated with lower multivariate decoding. However, whether and to what extent confidence judgements about internal states reflect a readout of discriminability between neural patterns remains an unresolved issue. Recent work demonstrated that confidence judgements and behavioural accuracy are dissociated during decision making and these dissociations can be explained by differences in neural computations (Peters et al., 2017). One limitation in this study is that due to the small number of trials per certain classes in certain participants, our analyses were limited to mean confidence ratings. Future research on mind wandering could utilize confidence ratings on a trial-by-trial basis to provide a more precise estimate of the relationship between decoding accuracy and classification accuracy (Weaver et al. 2019).

## Conclusions

Our findings expand upon research linking elevated alpha power with mind wandering episodes and reveal distinct electrophysiological characteristics of state meta-awareness. Unaware mind wandering was consistently more dissimilar from on-task states than aware mind wandering, as evidenced by superior decoding and greater neurophysiological differences. These results highlight a clear distinction between unaware and aware mind wandering states and confirm the utility of introspective methods in the study of transient fluctuations in conscious experience. The observed effects demonstrate the potential of using EEG machine learning classifiers to capture mind wandering and state meta-awareness during an ecological task without performance indicators. We found that confidence in experiential state and state meta-awareness reports correlated with the decoding of the respective states, suggesting that introspection confidence scales with neurophysiological dissimilarity. These effects suggest that introspection confidence taps into variability in metacognitive access to, and differential phenomenological characteristics, of experiential states.

## Data Availability

The data that support the findings of this study are available from the corresponding author upon request.

## Acknowledgments

MHR was partially funded by the Basic Research Program of the National Research University Higher School of Economics (Russian Federation). The authors would like to thank Stefan Haufe for engaging in useful discussions about activation patterns of MVPC analyses.

## References

1. Allefeld, C., Görgen, K., & Haynes, J.-D. (2016). Valid population inference for information-based imaging: From the second-level t-test to prevalence inference. NeuroImage, 141, 378–392. https://doi.org/10.1016/j.neuroimage.2016.07.040

2. Allefeld, C., & Haynes, J.-D. (2015). Multi-voxel Pattern Analysis. In Brain Mapping: An Encyclopedic Reference (Vol. 1). Elsevier Inc. https://doi.org/10.1016/B978-0-12-397025-1.00345-6

3. Andrews-Hanna, J. R. (2012). The Brain’s Default Network and Its Adaptive Role in Internal Mentation. The Neuroscientist, 18(3), 251–270. https://doi.org/10.1177/1073858411403316

4. Andrews-Hanna, J. R., Smallwood, J., & Spreng, R. N. (2014). The default network and self-generated thought: Component processes, dynamic control, and clinical relevance. Annals of the New York Academy of Sciences, 1316(1), 29–52. https://doi.org/10.1111/nyas.12360

5. Antrobus, J. S., Singer, J. L., Goldstein, S., & Fortgang, M. (1970). Section of Psychology: Mind Wandering and Cognitive Structure. Transactions of the New York Academy of Sciences, 32(2 Series II), 242–252. https://doi.org/10.1111/j.2164-0947.1970.tb02056.x

6. Bae, G. Y., & Luck, S. J. (2018). Dissociable decoding of spatial attention and working memory from EEG oscillations and sustained potentials. Journal of Neuroscience, 38(2), 409–422. https://doi.org/10.1523/JNEUROSCI.2860-17.2017

7. Baird, B., Smallwood, J., Lutz, A., & Schooler, J. W. (2014). The Decoupled Mind: Mind-wandering Disrupts Cortical Phase-locking to Perceptual Events. Journal of Cognitive Neuroscience, 26(11), 2596–2607. https://doi.org/10.1162/jocn_a_00656

8. Baldwin, C. L., Roberts, D. M., Barragan, D., Lee, J. D., Lerner, N., & Higgins, J. S. (2017). Detecting and quantifying mind wandering during simulated driving. Frontiers in Human Neuroscience, 11(August), 1–15. https://doi.org/10.3389/fnhum.2017.00406

9. Barron, E., Riby, L. M., Greer, J., & Smallwood, J. (2011). Absorbed in Thought. Psychological Science, 22(5), 596–601. https://doi.org/10.1177/0956797611404083

10. Benjamini, Y., Krieger, A. M., & Yekutieli, D. (2006). Adaptive linear step-up procedures that control the false discovery rate. Biometrika, 93(3), 491–507. https://doi.org/10.1093/biomet/93.3.491

11. Boldt, A., & Yeung, N. (2015). Shared neural markers of decision confidence and error detection. Journal of Neuroscience, 35(8), 3478–3484. https://doi.org/10.1523/JNEUROSCI.0797-14.2015

12. Boudewyn, M. A., & Carter, C. S. (2018). I must have missed that: Alpha-band oscillations track attention to spoken language. Neuropsychologia, 117(April), 148–155. https://doi.org/10.1016/j.neuropsychologia.2018.05.024

13. Braboszcz, C., & Delorme, A. (2011). Lost in thoughts: Neural markers of low alertness during mind wandering. NeuroImage, 54(4), 3040–3047. https://doi.org/10.1016/j.neuroimage.2010.10.008

14. Broadway, J. M., Franklin, M. S., & Schooler, J. W. (2015). Early event-related brain potentials and hemispheric asymmetries reveal mind-wandering while reading and predict comprehension. Biological Psychology, 107, 31–43. https://doi.org/10.1016/j.biopsycho.2015.02.009

15. Bryson, B. (2004). A short history of nearly everything. Black Swan.

16. Cavanagh & Frank, M. J. (2014). Frontal Theta as a Mechanism for Affective and Effective Control. Trends in Cognitive Sciences, 18(8), 414–421. https://doi.org/10.1016/j.tics.2014.04.012.Frontal

17. Chang, C. C., & Lin, C. J. (2011). LIBSVM: A Library for support vector machines. ACM Transactions on Intelligent Systems and Technology, 2, 1–27. https://doi.org/10.1145/1961189.1961199

18. Christoff, K., Gordon, A. M., Smallwood, J., Smith, R., & Schooler, J. W. (2009). Experience sampling during fMRI reveals default network and executive system contributions to mind wandering. Proceedings of the National Academy of Sciences of the United States of America, 106(21), 8719–8724. https://doi.org/10.1073/pnas.0900234106

19. Cohen, M. X. (2014). A neural microcircuit for cognitive conflict detection and signaling. Trends in Neurosciences, 37(9), 480–490. https://doi.org/10.1016/j.tins.2014.06.004

20. Compton, R. J., Gearinger, D., & Wild, H. (2019). The wandering mind oscillates: EEG alpha power is enhanced during moments of mind-wandering. *Cognitive*, Affective and Behavioral Neuroscience, 19(5), 1184–1191. https://doi.org/10.3758/s13415-019-00745-9

21. Cooper, N. R., Croft, R. J., Dominey, S. J. J., Burgess, A. P., & Gruzelier, J. H. (2003). Paradox lost? Exploring the role of alpha oscillations during externally vs. internally directed attention and the implications for idling and inhibition hypotheses. International Journal of Psychophysiology, 47(1), 65–74. https://doi.org/10.1016/S0167-8760(02)00107-1

22. Delorme, A., & Makeig, S. (2004). EEGLAB: An open source toolbox for analysis of single-trial EEG dynamics including independent component analysis. Journal of Neuroscience Methods, 134(1), 9–21. https://doi.org/10.1016/j.jneumeth.2003.10.009

23. DeLosAngeles, D., Williams, G., Burston, J., Fitzgibbon, S. P., Lewis, T. W., Grummett, T. S., Clark, C. R., Pope, K. J., & Willoughby, J. O. (2016). Electroencephalographic correlates of states of concentrative meditation. International Journal of Psychophysiology, 110(October 2017), 27–39. https://doi.org/10.1016/j.ijpsycho.2016.09.020

24. Dhindsa, K., Acai, A., Wagner, N., Bosynak, D., Kelly, S., Bhandari, M., Petrisor, B., & Sonnadara, R. R. (2019). Individualized pattern recognition for detecting mind wandering from EEG during live lectures. PLoS ONE, 14(9), 1–30. https://doi.org/10.1371/journal.pone.0222276

25. Efron, B. (1987). Better bootstrap confidence intervals. Journal of the American Statistical Association, 82(397), 171–185. https://doi.org/10.1080/01621459.1987.10478410

26. Feng, S., D’Mello, S., & Graesser, A. C. (2013). Mind wandering while reading easy and difficult texts. Psychonomic Bulletin & Review, 20(3), 586–592. https://doi.org/10.3758/s13423-012-0367-y

27. Fingelkurts, A. A., & Fingelkurts, A. A. (2011). Persistent operational synchrony within brain default-mode network and self-processing operations in healthy subjects. Brain and Cognition, 75(2), 79–90. https://doi.org/10.1016/j.bandc.2010.11.015

28. Fleming, S. M., & Lau, H. C. (2014). How to measure metacognition. Frontiers in Human Neuroscience, 8, 443. https://doi.org/10.3389/fnhum.2014.00443

29. Fox, M. D., Snyder, A. Z., Vincent, J. L., Corbetta, M., Van Essen, D. C., & Raichle, M. E. (2005). The human brain is intrinsically organized into dynamic, anticorrelated functional networks. Proceedings of the National Academy of Sciences of the United States of America, 102(27), 9673–9678. https://doi.org/10.1073/pnas.0504136102

30. Foxe, J. J., & Snyder, A. C. (2011). The role of alpha-band brain oscillations as a sensory suppression mechanism during selective attention. Frontiers in Psychology, 2(JUL), 1–13. https://doi.org/10.3389/fpsyg.2011.00154

31. Geden, M., Staicu, A.-M., & Feng, J. (2018). The impacts of perceptual load and driving duration on mind wandering in driving. Transportation Research Part F: Traffic Psychology and Behaviour, 57, 75–83. https://doi.org/10.1016/j.trf.2017.07.004

32. Gherman, S., & Philiastides, M. G. (2018). Human VMPFC encodes early signatures of confidence in perceptual decisions. ELife, 7, 1–28. https://doi.org/10.7554/eLife.38293

33. Grandchamp, R., & Delorme, A. (2011). Single-trial normalization for event-related spectral decomposition reduces sensitivity to noisy trials. Frontiers in Psychology, 2(SEP), 1–14. https://doi.org/10.3389/fpsyg.2011.00236

34. Groot, J. M., Boayue, N. M., Csifcsák, G., Boekel, W., Huster, R., Forstmann, B. U., & Mittner, M. (2021). Probing the neural signature of mind wandering with simultaneous fMRI-EEG and pupillometry. NeuroImage, 224(June 2020). https://doi.org/10.1016/j.neuroimage.2020.117412

35. Hanslmayr, S., Gross, J., Klimesch, W., & Shapiro, K. L. (2011). The role of alpha oscillations in temporal attention. Brain Research Reviews, 67(1–2), 331–343. https://doi.org/10.1016/j.brainresrev.2011.04.002

36. Harmony, T. (2013). The functional significance of delta oscillations in cognitive processing. Frontiers in Integrative Neuroscience, 7(DEC), 1–10. https://doi.org/10.3389/fnint.2013.00083

37. Harmony, T., Fernández, T., Silva, J., Bernal, J., Díaz-Comas, L., Reyes, A., Marosi, E., Rodríguez, M., & Rodríguez, M. (1996). EEG delta activity: An indicator of attention to internal processing during performance of mental tasks. International Journal of Psychophysiology, 24(1–2), 161–171. https://doi.org/10.1016/S0167-8760(96)00053-0

38. Haufe, S., Meinecke, F., Görgen, K., Dähne, S., Haynes, J. D., Blankertz, B., & Bießmann, F. (2014). On the interpretation of weight vectors of linear models in multivariate neuroimaging. NeuroImage, 87(November), 96–110. https://doi.org/10.1016/j.neuroimage.2013.10.067

39. Haxby, J. V., Gobbini, M. I., Furey, M. L., Ishai, A., Schouten, J. L., & Pietrini, P. (2001). Distributed and overlapping representations of faces and objects in ventral temporal cortex. Science, 293(5539), 2425–2430. https://doi.org/10.1126/science.1063736

40. Haynes, J. D., & Rees, G. (2006). Decoding mental states from brain activity in humans. Nature Reviews Neuroscience, 7(7), 523–534. https://doi.org/10.1038/nrn1931

41. Hollis, R. B., & Was, C. A. (2016). Mind wandering, control failures, and social media distractions in online learning. Learning and Instruction, 42, 104–112. https://doi.org/10.1016/j.learninstruc.2016.01.007

42. Jin, C. Y., Borst, J. P., & van Vugt, M. K. (2019). Predicting task-general mind-wandering with EEG. *Cognitive*, Affective and Behavioral Neuroscience, 19(4), 1059–1073. https://doi.org/10.3758/s13415-019-00707-1

43. Kam, J. W. Y., Dao, E., Farley, J., Fitzpatrick, K., Smallwood, J., Schooler, J. W., & Handy, T. C. (2011). Slow Fluctuations in Attentional Control of Sensory Cortex. Journal of Cognitive Neuroscience, 23(2), 460–470. https://doi.org/10.1162/jocn.2010.21443

44. Killingsworth, M. A., & Gilbert, D. T. (2010). A wandering mind is an unhappy mind. Science, 330(6006), 932. https://doi.org/10.1126/science.1192439

45. Kim, J., Chung, Y. G., Park, J. Y., Chung, S. C., Wallraven, C., Bülthoff, H. H., & Kim, S. P. (2015). Decoding accuracy in supplementary motor cortex correlates with perceptual sensitivity to tactile roughness. PLoS ONE, 10(6), 1–17. https://doi.org/10.1371/journal.pone.0129777

46. Kleiner, M., Brainard, D., Pelli, D., Ingling, A., Murray, R., & Broussard, C. (2007). Perception. In Perception (Vol. 36, Issue 14). [Pion Ltd.]. https://nyuscholars.nyu.edu/en/publications/whats-new-in-psychtoolbox-3

47. Klimesch, W. (1999). EEG alpha and theta oscillations reflect cognitive and memory performance: a rKlimesch, W. (1999). EEG alpha and theta oscillations reflect cognitive and memory performance: a review and analysis. Brain Research Reviews, 29(2-3), 169–195. doi:10.1016/S016. *Brain Research Reviews*, *29*(2–3), 169–195. https://doi.org/10.1016/S0165-0173(98)00056-3

48. Klimesch, W. (2012). Alpha-band oscillations, attention, and controlled access to stored information. Trends in Cognitive Sciences, 16(12), 606–617. https://doi.org/10.1016/j.tics.2012.10.007

49. Kubanek, J., Hill, N. J., Snyder, L. H., & Schalk, G. (2015). Cortical alpha activity predicts the confidence in an impending action. Frontiers in Neuroscience, 9(JUL), 1–15. https://doi.org/10.3389/fnins.2015.00243

50. Kunimoto, C., Miller, J., & Pashler, H. (2001). Confidence and accuracy of near-threshold discrimination responses. Consciousness and Cognition, 10(3), 294–340. https://doi.org/10.1006/ccog.2000.0494

51. Lindquist, S. I., & McLean, J. P. (2011). Daydreaming and its correlates in an educational environment. Learning and Individual Differences, 21(2), 158–167. https://doi.org/10.1016/j.lindif.2010.12.006

52. Lotte, F., Congedo, M., Lécuyer, A., Lamarche, F., & Arnaldi, B. (2007). A review of classification algorithms for EEG-based brain-computer interfaces. Journal of Neural Engineering, 4(2). https://doi.org/10.1088/1741-2560/4/2/R01

53. Macdonald, J. S. P., Mathan, S., & Yeung, N. (2011). Trial-by-trial variations in subjective attentional state are reflected in ongoing prestimulus EEG alpha oscillations. Frontiers in Psychology, 2, 82. https://doi.org/10.3389/fpsyg.2011.00082

54. Maris, E., & Oostenveld, R. (2007). Nonparametric statistical testing of EEG- and MEG-data. Journal of Neuroscience Methods, 164(1), 177–190. https://doi.org/10.1016/j.jneumeth.2007.03.024

55. Martinon, L. M., Smallwood, J., McGann, D., Hamilton, C., & Riby, L. M. (2019). The disentanglement of the neural and experiential complexity of self-generated thoughts: A users guide to combining experience sampling with neuroimaging data. NeuroImage, 192(September 2018), 15–25. https://doi.org/10.1016/j.neuroimage.2019.02.034

56. McVay, J. C., Kane, M. J., & Kwapil, T. R. (2009). Tracking the train of thought from the laboratory into everyday life: An experience-sampling study of mind wandering across controlled and ecological contexts. Psychonomic Bulletin & Review, 16(5), 857–863. https://doi.org/10.3758/PBR.16.5.857

57. Meier, M. E. (2018). Can research participants comment authoritatively on the validity of their self-reports of mind wandering and task engagement? A replication and extension of Seli, Jonker, Cheyne, Cortes, and Smilek (2015). Journal of Experimental Psychology: Human Perception and Performance, 44(10), 1567–1585. https://doi.org/10.1037/xhp0000556

58. Mitchell, D. J., McNaughton, N., Flanagan, D., & Kirk, I. J. (2008). Frontal-midline theta from the perspective of hippocampal “theta.” Progress in Neurobiology, 86(3), 156–185. https://doi.org/10.1016/j.pneurobio.2008.09.005

59. Mittner, M., Boekel, W., Tucker, A. M., Turner, B. M., Heathcote, A., & Forstmann, B. U. (2014). When the brain takes a break: A model-based analysis of mind wandering. Journal of Neuroscience, 34(49), 16286–16295. https://doi.org/10.1523/JNEUROSCI.2062-14.2014

60. Morgan, M. J., Mason, A. J. S., & Solomon, J. A. (1997). Blindsight in normal subjects? [10]. Nature, *385*(6615), 401–402. https://doi.org/10.1038/385401b0

61. Palva, J. M., Palva, S., & Kaila, K. (2005). Phase synchrony among neuronal oscillations in the human cortex. Journal of Neuroscience, 25(15), 3962–3972. https://doi.org/10.1523/JNEUROSCI.4250-04.2005

62. Palva, S., & Palva, J. M. (2007). New vistas for α-frequency band oscillations. Trends in Neurosciences, 30(4), 150–158. https://doi.org/10.1016/j.tins.2007.02.001

63. Pernet, C. R., Wilcox, R., & Rousselet, G. A. (2013). Robust correlation analyses: False positive and power validation using a new open source matlab toolbox. Frontiers in Psychology. https://doi.org/10.3389/fpsyg.2012.00606

64. Peters, M. A. K., Thesen, T., Ko, Y. D., Maniscalco, B., Carlson, C., Davidson, M., Doyle, W., Kuzniecky, R., Devinsky, O., Halgren, E., & Lau, H. (2017). Perceptual confidence neglects decision-incongruent evidence in the brain. Nature Human Behaviour, 1(7), 1–8. https://doi.org/10.1038/s41562-017-0139

65. Raichle, M. E., MacLeod, A. M., Snyder, A. Z., Powers, W. J., Gusnard, D. A., & Shulman, G. L. (2001). A default mode of brain function. Proceedings of the National Academy of Sciences of the United States of America, 98(2), 676–682. https://doi.org/10.1073/pnas.98.2.676

66. Rousseeuw, P. J. (1984). Least median of squares regression. Journal of the American Statistical Association, 79(388), 871–880. https://doi.org/10.1080/01621459.1984.10477105

67. Rousseeuw, P. J., & Van Driessen, K. (1999). A fast algorithm for the minimum covariance determinant estimator. Technometrics, 41(3), 212–223. https://doi.org/10.1080/00401706.1999.10485670

68. Sauseng, P., Klimesch, W., Doppelmayr, M., Pecherstorfer, T., Freunberger, R., & Hanslmayr, S. (2005). EEG alpha synchronization and functional coupling during top-down processing in a working memory task. Human Brain Mapping, 26(2), 148–155. https://doi.org/10.1002/hbm.20150

69. Schooler, J. W., Reichle, E. D., & Halpern, D. V. (2004). Zoning Out while Reading: Evidence for Dissociations between Experience and Metaconsciousness. In Thinking and seeing: Visual metacognition in adults and children (pp. 203–226). MIT Press.

70. Seli, P., Jonker, T. R., Cheyne, J. A., Cortes, K., & Smilek, D. (2015). Can research participants comment authoritatively on the validity of their self-reports of mind wandering and task engagement? Journal of Experimental Psychology: Human Perception and Performance, 41(3), 703–709. https://doi.org/10.1037/xhp0000029

71. Seli, P., Ralph, B. C. W., Risko, E. F., W. Schooler, J., Schacter, D. L., & Smilek, D. (2017). Intentionality and meta-awareness of mind wandering: Are they one and the same, or distinct dimensions? Psychonomic Bulletin and Review, 24(6), 1808–1818. https://doi.org/10.3758/s13423-017-1249-0

72. Seli, P., Risko, E. F., Smilek, D., & Schacter, D. L. (2016). Mind-Wandering With and Without Intention. Trends in Cognitive Sciences, 20(8), 605–617. https://doi.org/10.1016/j.tics.2016.05.010

73. Smallwood, J., Beach, E., Schooler, J. W., & Handy, T. C. (2008). Going AWOL in the Brain: Mind Wandering Reduces Cortical Analysis of External Events. Journal of Cognitive Neuroscience, 20(3), 458–469. https://doi.org/10.1162/jocn.2008.20037

74. Smallwood, J., Brown, K., Baird, B., & Schooler, J. W. (2012). Cooperation between the default mode network and the frontal-parietal network in the production of an internal train of thought. Brain Research, 1428, 60–70. https://doi.org/10.1016/j.brainres.2011.03.072

75. Smallwood, J., Fishman, D. J., & Schooler, J. W. (2007). Counting the cost of an absent mind: Mind wandering as an underrecognized influence on educational performance. Psychonomic Bulletin & Review, 14(2), 230–236. https://doi.org/10.3758/BF03194057

76. Smallwood, J., McSpadden, M., & Schooler, J. W. (2007). The lights are on but no one’s home: Meta-awareness and the decoupling of attention when the mind wanders. Psychonomic Bulletin and Review, 14(3), 527–533. https://doi.org/10.3758/BF03194102

77. Smallwood, J., Nind, L., & O’Connor, R. C. (2009). When is your head at? An exploration of the factors associated with the temporal focus of the wandering mind. Consciousness and Cognition, 18(1), 118–125. https://doi.org/10.1016/j.concog.2008.11.004

78. Smallwood, J., & Schooler, J. W. (2006). The restless mind. Psychological Bulletin, 132(6), 946– 958. https://doi.org/10.1037/0033-2909.132.6.946

79. Smallwood, J., & Schooler, J. W. (2015). The Science of Mind Wandering: Empirically Navigating the Stream of Consciousness. Annual Review of Psychology, 66(1), 487–518. https://doi.org/10.1146/annurev-psych-010814-015331

80. Spreng, R. N., & Grady, C. L. (2010). Patterns of brain activity supporting autobiographical memory, prospection, and theory of mind, and their relationship to the default mode network. Journal of Cognitive Neuroscience, 22(6), 1112–1123. https://doi.org/10.1162/jocn.2009.21282

81. Thielen, J., Bosch, S. E., van Leeuwen, T. M., van Gerven, M. A. J., & van Lier, R. (2019). Evidence for confounding eye movements under attempted fixation and active viewing in cognitive neuroscience. Scientific Reports, 9(1), 1–8. https://doi.org/10.1038/s41598-019-54018-z

82. Vago, D. R., & Zeidan, F. (2016). The brain on silent: mind wandering, mindful awareness, and states of mental tranquility. Annals of the New York Academy of Sciences, 1373(1), 96–113. https://doi.org/10.1111/nyas.13171

83. van Son, D., De Blasio, F. M., Fogarty, J. S., Angelidis, A., Barry, R. J., & Putman, P. (2019). Frontal EEG theta/beta ratio during mind wandering episodes. Biological Psychology, 140(November 2018), 19–27. https://doi.org/10.1016/j.biopsycho.2018.11.003

84. Varao Sousa, T. L., Carriere, J. S. A., & Smilek, D. (2013). The way we encounter reading material influences how frequently we mind wander. Frontiers in Psychology, 4, 892. https://doi.org/10.3389/fpsyg.2013.00892

85. Verboven, S., & Hubert, M. (2005). LIBRA: A MATLAB library for robust analysis. Chemometrics and Intelligent Laboratory Systems, 75(2), 127–136. https://doi.org/10.1016/j.chemolab.2004.06.003

86. Villena-González, M., López, V., & Rodríguez, E. (2016). Orienting attention to visual or verbal/auditory imagery differentially impairs the processing of visual stimuli. NeuroImage, 132, 71–78. https://doi.org/10.1016/j.neuroimage.2016.02.013

87. Von Stein, A., & Sarnthein, J. (2000). Different frequencies for different scales of cortical integration: From local gamma to long range alpha/theta synchronization. International Journal of Psychophysiology, 38(3), 301–313. https://doi.org/10.1016/S0167-8760(00)00172-0

88. Weaver, M. D., Fahrenfort, J. J., Belopolsky, A., & Van Gaal, S. (2019). Independent neural activity patterns for sensory-and confidence-based information maintenance during category-selective visual processing. ENeuro, 6(1). https://doi.org/10.1523/ENEURO.0268-18.2018

89. Wokke, M. E., Cleeremans, A., & Ridderinkhof, K. R. (2017). Sure i’m sure: Prefrontal oscillations support metacognitive monitoring of decision making. Journal of Neuroscience, 37(4), 781– 789. https://doi.org/10.1523/JNEUROSCI.1612-16.2016

90. Zakrzewski, A. C., Wisniewski, M. G., Iyer, N., & Simpson, B. D. (2019). Confidence tracks sensory- and decision-related ERP dynamics during auditory detection. Brain and Cognition, 129(October 2018), 49–58. https://doi.org/10.1016/j.bandc.2018.10.007

