## Supplementary Materials for "Introspection confidence predicts EEG decoding of self-generated thoughts and meta-awareness"

**Supplementary Results: Cluster-based permutations**

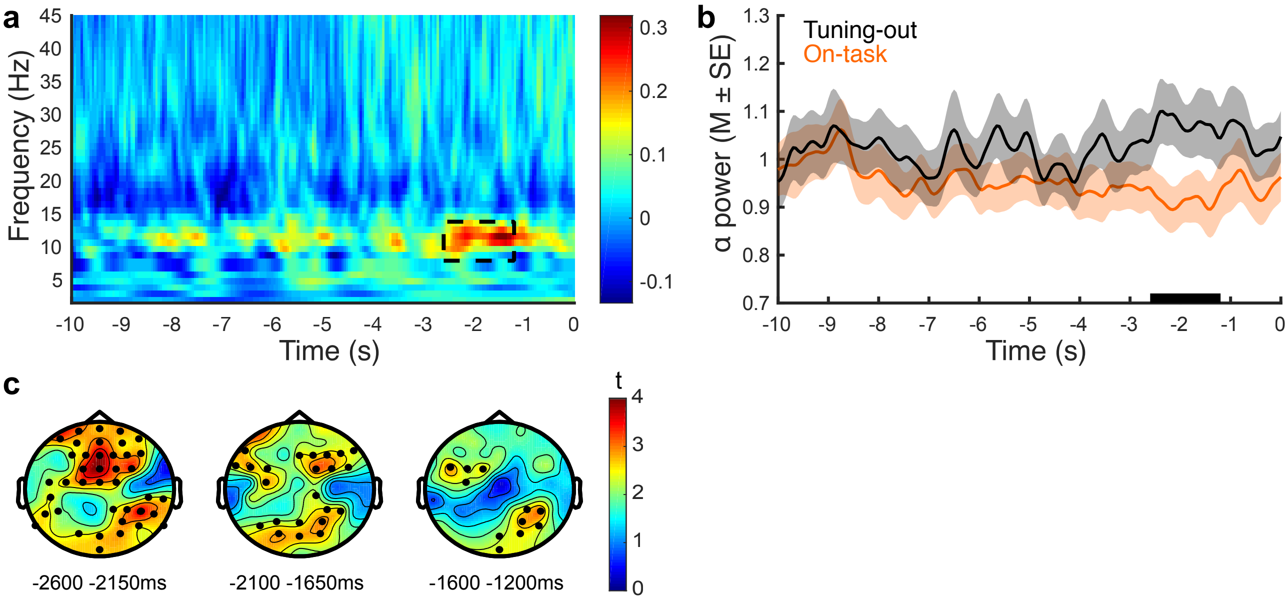


**Supplementary Figure 1.** Oscillatory differences between states (tuning-out – on-task, *N*=27) as a function of time relative to probe onset (0s). a) Time frequency decomposition averaged across all electrode sites. The broken black rectangle indicates the spectrotemporal cluster reflecting significant state differences ($p<.025$, two-sided cluster-based permutation test). b) Alpha (8-13Hz) spectral power averaged over the electrode sites of the cluster (significance denoted by a black bar on the x-axis). c) Topography of the cluster at different 400/450ms sub-windows (black markers denote electrodes that were present in at least 50% of samples in each time window).

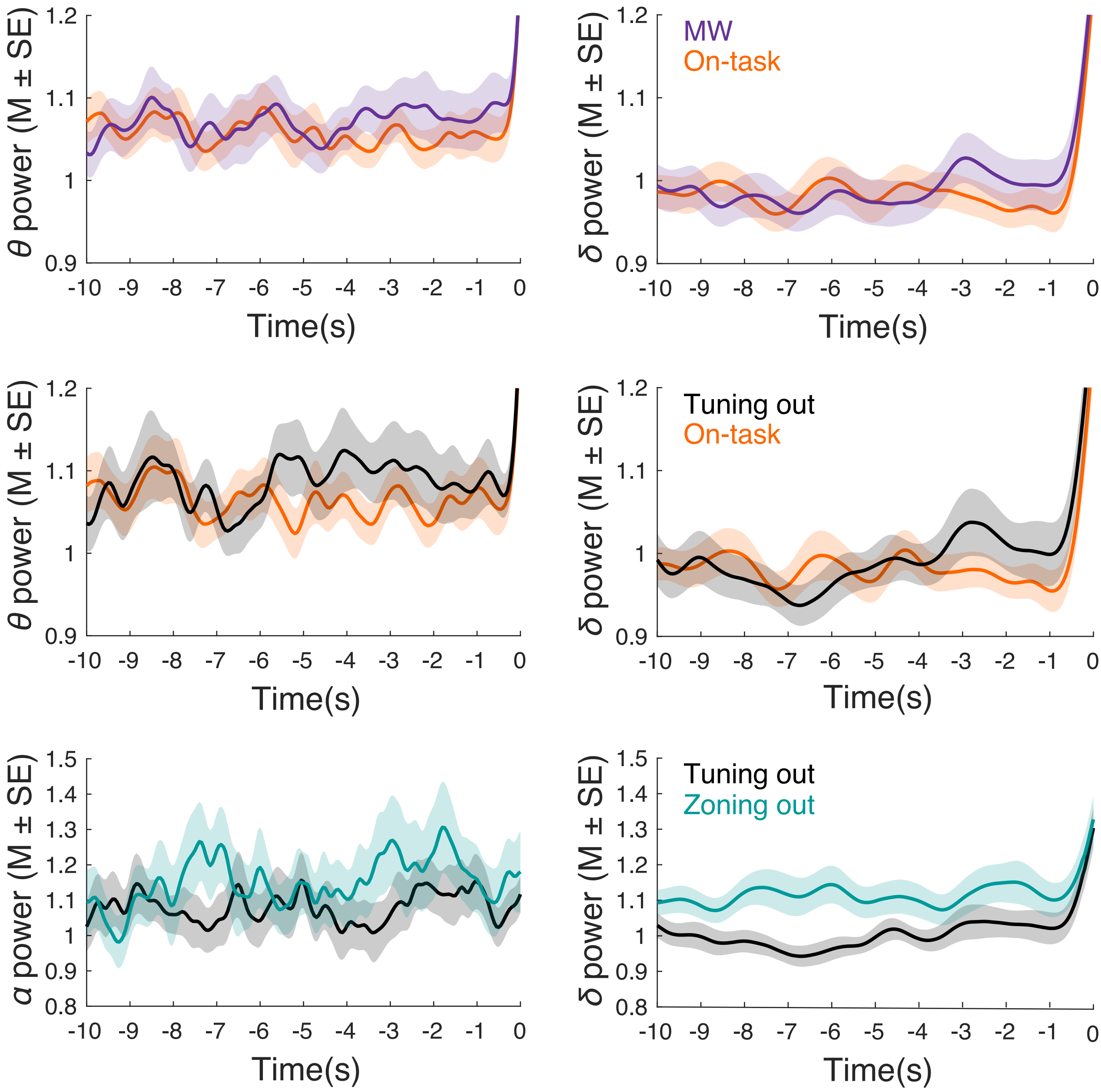


**Supplementary Figure 2.** Spectral power averaged over all 64 electrode sites as a function of frequency band and condition for contrasts that were non-significant; We report the strongest, albeit non-significant, effects. Top panel: the on-task vs. MW analysis did not yield any significant clusters for delta ($p$=.24, at -4.500 to -3.750s) or theta power ($p$=.40, at -3.550 to -3.200, negative cluster). Middle panel: theta and delta power also did not significantly differ between tuning-out and on-task states (theta: $p$=.20, at -7.500 to -7.050s, delta: $p$=.23 at -8.250 to -7.750s, negative cluster). Bottom panel: tuning-out and zoning-out states did not differ in alpha ($p$=.16, at -7.500 to 6.850s) or delta power ($p$=.05, at -8.400 to -7.050). MW = Mind Wandering.

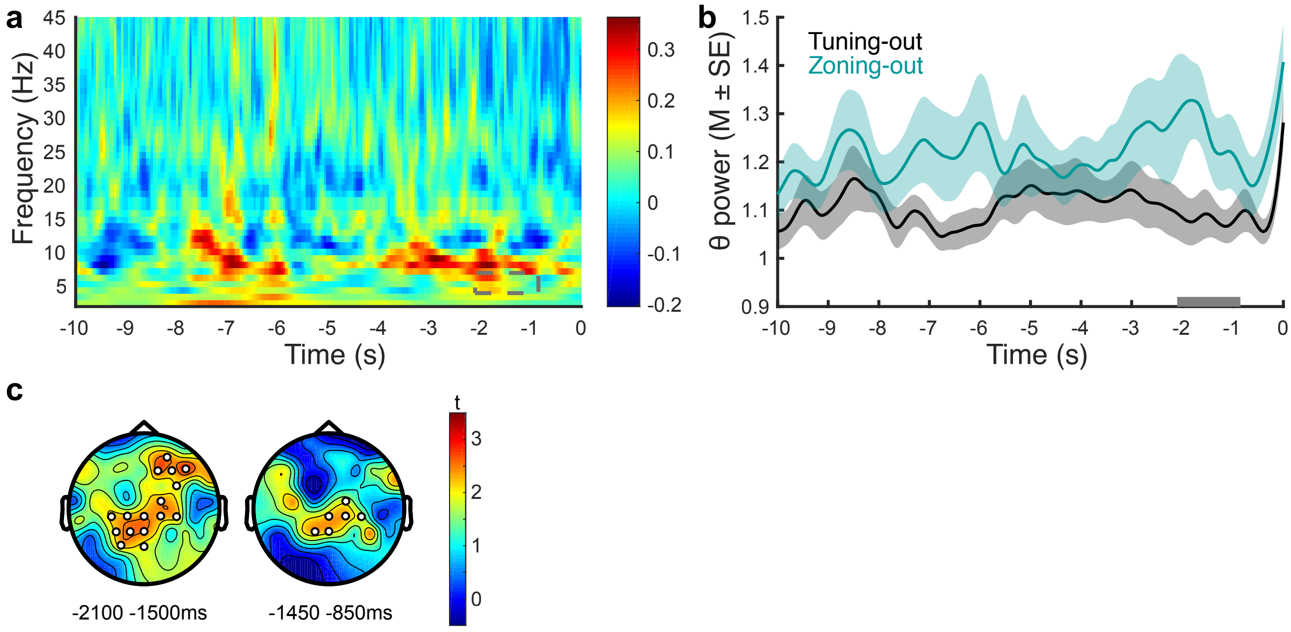


**Supplementary Figure 3.** Oscillatory differences between states (zoning-out - tuning-out, *N*=21) as a function of time relative to probe onset (0s). a) Time frequency decomposition averaged across electrode sites. The broken grey rectangle denotes a spectrotemporal cluster reflecting the largest identified state differences (*p*=.026, two-sided cluster-based permutation test). b) Theta (4-7Hz) spectral power averaged over the electrode sites of the cluster (trend effect denoted by grey bar on the x-axis). c) Topography of the cluster at two 600ms sub-windows (white markers denote electrodes that were present at least 50% of samples in time window).

**Control analyses: Cluster-based permutations**

Insofar as the number of trials differed between states, this represents a potential confound in the state contrasts of oscillatory power. In order to address this, we conducted control analyses minimizing discrepancies between inter-condition trial numbers, while maintaining a good sample size using two criteria: we removed participants with the highest discrepancies until either the difference in mean trial numbers was smaller than 20%, or until we reached 60% of each sample. The four main contrasts were implemented with the following trial numbers ($M \pm SD$) and sample sizes: (i) on-task (35.2±9.8) vs. mind wandering (28.6±10.0) [$n$=29]; (ii) on-task (29.8±8.9) vs. tuning-out (23.2±8.6) [$n$=16]; (iii) on-task (27.9±9.8) vs. zoning-out (15.1±5.3) [$n$=15]; and (iv) tuning-out (16.7±5.4) vs. zoning-out (13.7±4.3) [$n$=17]. Cluster-based permutations analyses were conducted in the frequency bands that were significant or at a trend-level in the main analyses.

In line with the main findings, the analysis between mind wandering and on-task trials showed that mind wandering compared to on-task states was associated with greater alpha power close to probe onset (-3.800 to 1.100s), $p$=.016. Likewise, tuning-out was associated with greater alpha power than on-task states in one cluster at -2.900 to -1.200 seconds prior to probe onset, $p$=.004. Comparison between zoning-out and on-task states revealed greater alpha power within a large time window (-4.400 to -0.850s), $p$=.009. An earlier cluster (-7.900 to -6.700s) was comparable to the main analysis, albeit was non-significant, $p$=.050. Theta power was also greater for zoning-out compared to on-task states, $p$=.020 (-2.250 to -1.100s), although delta effects in this reduced sample were not significant, $p$=.068 (-7.950 to -6.950s). Finally, a comparison between zoning-out and tuning-out states showed that theta power was greater in zoning-out states; however, in line with the main analysis, this remained at trend-level, $p$=.026 (-2.250 to -0.950s). These results demonstrate that the significant differences between states reported in the main findings are unlikely to be an artifact of discrepancies due to trial numbers.

**Exploratory Analyses: Beta power cluster-based permutations**

On the basis of previous research which yielded preliminary evidence for a potential role of beta oscillations in mind wandering (Braboszcz & Delorme, 2011; van Son et al., 2019) we also examined whether there were any power differences between mind wandering and on-task states in the beta (14-30Hz) frequency band. These analyses did not yield any significant effects, therefore we report *p* values for the most prominent cluster in **Supplementary Figure 4**. As is evident, we failed to find any evidence of an association between beta power and mind wandering or state meta-awareness.

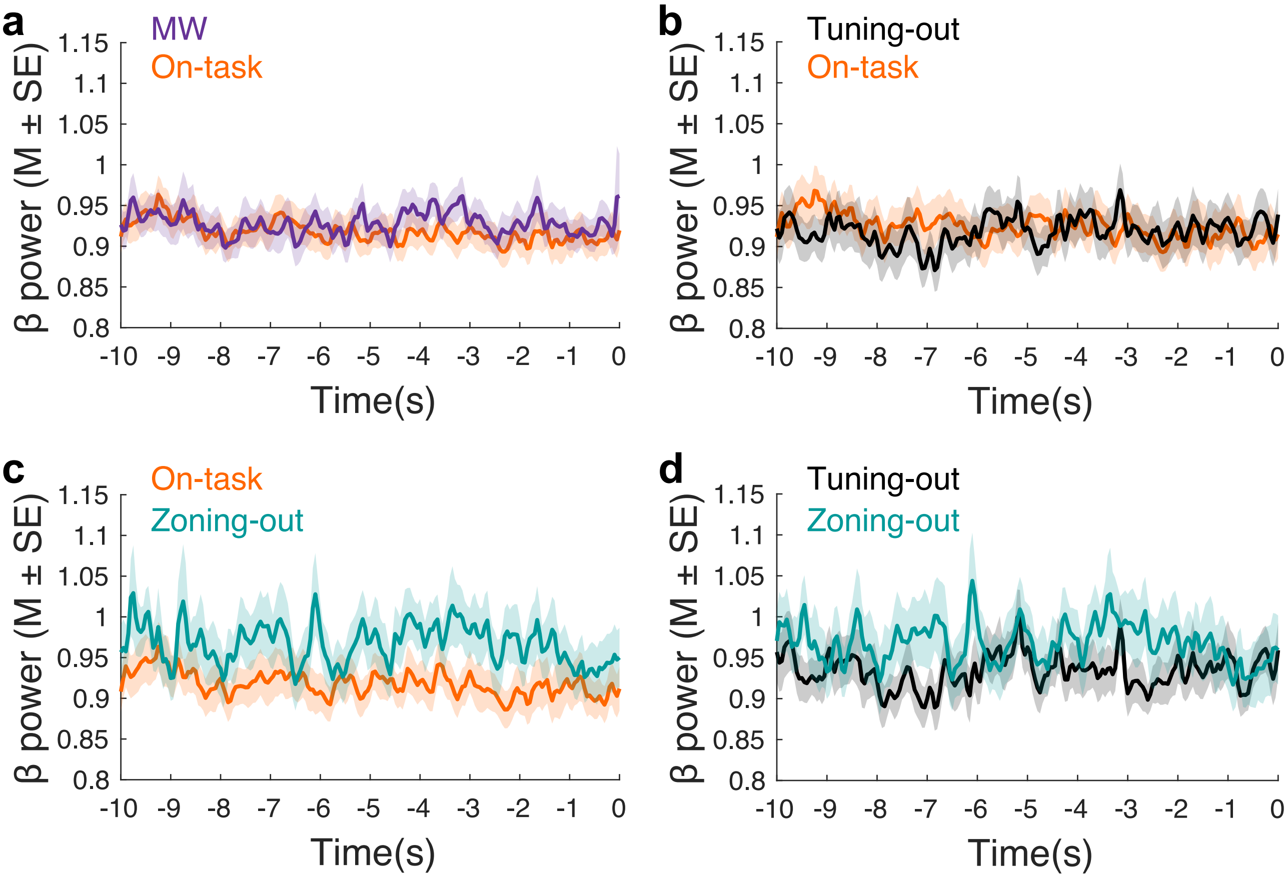


**Supplementary Figure 4.** Beta (14-30Hz) spectral power averaged over all 64 electrode sites: a) MW vs. on-task (*N*=39): *p*=.41 (at -4.400 to -4.300s), b) On-task vs. tuning-out (*N*=27): *p*=.35 (at -7.250 to -7.100), c) Zoning out vs. on-task (*N*=25): *p*=.16 (at -3.300 to -3.100), d) Zoning-out vs. tuning-out, (*N*=21): *p*=.15 (at 7.150 to -6.850). *P*-values correspond to the most prominent non-significant clusters. MW = Mind Wandering.

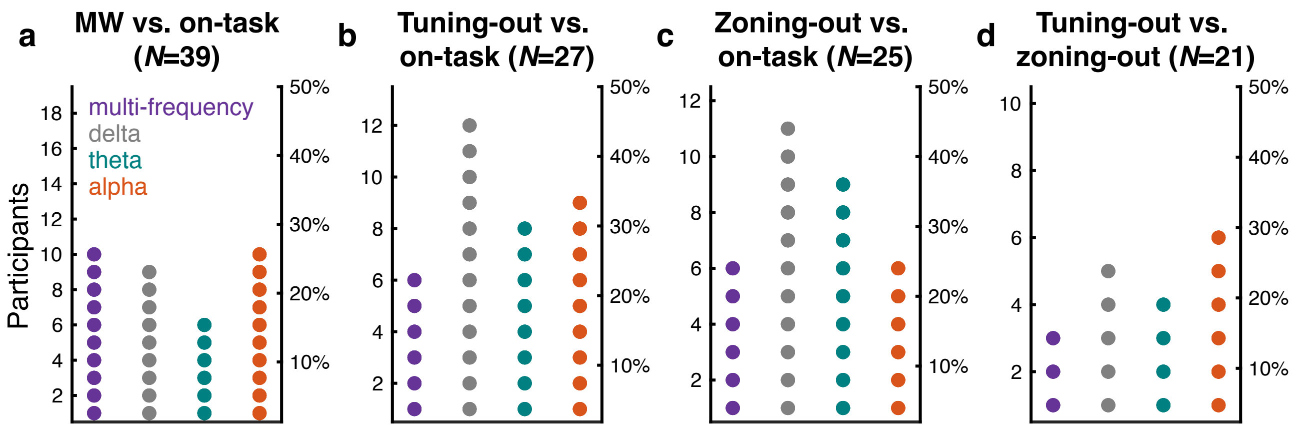


**Supplementary Figure 5.** Time-varying MVPC (Multivariate Pattern Classification analysis): Participants (count and percentage) with significant (*p*<*pth*) decoding in at least one time bin. MW = Mind Wandering.

**Supplementary Table 1**

*Time-varying MVPC (Multivariate Pattern Classification analysis) – Group-level analyses FDR (False Discovery Rate) threshold p-values*

| | Multi-frequency | Delta | Theta | Alpha |
|---|---|---|---|---|
| MW vs. on-task ($N$=39) | $p \leq pth$=.002 | $p \leq pth$=.026 | $p \leq pth$ =.018 | $p \leq pth$ =.008 |
| Tuning-out vs. on-task ($N$=27) | $p \leq pth$ =.016 | ns, $p > pth$ | $p \leq pth$ =.017 | $p \leq pth$ =.005 |
| Zoning-out vs. on-task ($N$=25) | $p \leq pth$=.003 | $p \leq pth$ =.300 | $p \leq pth$=.037 | $p \leq pth$ <.001 |
| Zoning-out vs. tuning-out ($N$=21) | $p \leq pth$<.001 | $p \leq pth$=.016 | $p \leq pth$=.021 | $p \leq pth$ =.006 |

* These correspond to Figure 3 of the main text.

* MW = Mind Wandering.

**Supplementary Results: Time-averaged MVPC**

Time-averaged MVPC was complemented with a method that allows assessing the activation patterns that describe the contribution of each channel to the decoding of class-related information (Haufe et al., 2014). For decoding tuning-out and on-task states (**Supplementary Figure 6**) analysis showed that, for the multi-frequency and alpha models, class-related information was strongly present at fronto-central electrodes and was associated with negative values. In the multi-frequency model class-related information was additionally evident at posterior and frontal sites associated with positive values. In the theta model, class-related information was strongly present at posterior sites across participants and was associated with positive values. In the delta model, positive weight values of larger amplitude were observed at centro-parietal and posterior sites.

For zoning-out vs. on-task states (**Supplementary Figure 6**), the larger amplitude changes in activation maps were associated with negative values for the multi-frequency, theta, and alpha models at central, right frontal and right central electrodes respectively, suggesting that these sites represented class-related information more strongly. By contrast, in the delta model, frontal as well as posterior electrodes represented class-related information more strongly, through positive activation values.

For decoding state meta-awareness (tuning-out vs. zoning-out, **Supplementary Figure 6**), negative activation values were observed for the multi-frequency and theta models at right frontal regions, suggesting these regions represented class-related information more strongly (with a negative direction of the effect). In the delta and alpha models, class-related information was strongly present at central electrodes, in association with negative values. Additionally, in the delta model, class-related information was strongly present at fronto-central electrodes in association with positive values.

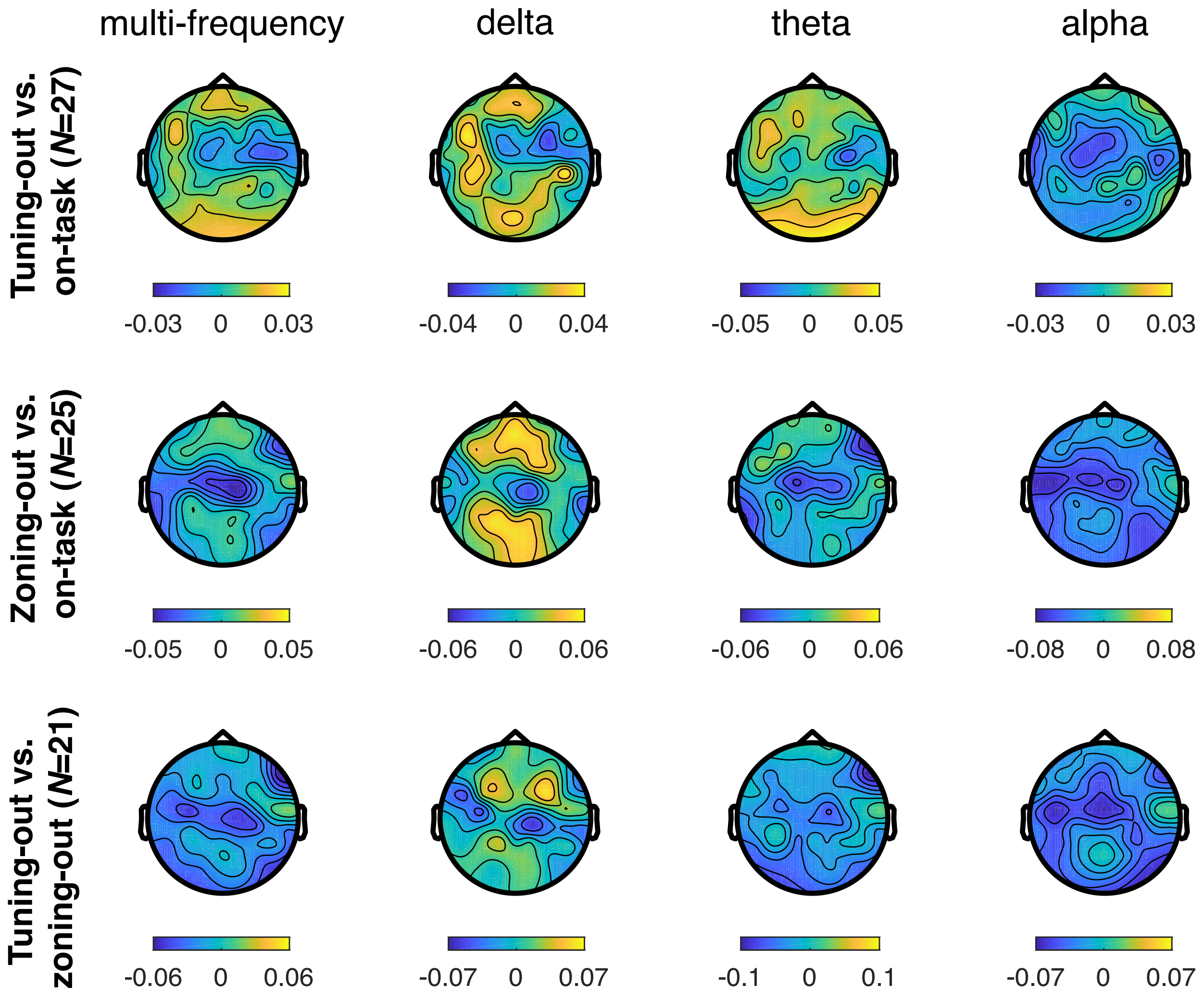


**Supplementary Figure 6.** Activation patterns indicative of the strength and polarity with which class-related information is reflected in each pattern dimension (electrode site) for the time-averaged MVPC (Multivariate Pattern Classification analysis). Each column displays activation patterns for each model (multi-frequency, delta, theta, alpha), whereas each row corresponds to a different two-class comparison. Activation values have arbitrary units.

### References


Braboszcz, C., & Delorme, A. (2011). Lost in thoughts: Neural markers of low alertness during mind wandering. *NeuroImage*, *54*(4), 3040–3047. https://doi.org/10.1016/j.neuroimage.2010.10.008

Haufe, S., Meinecke, F., Görgen, K., Dähne, S., Haynes, J. D., Blankertz, B., & Bießmann, F. (2014). On the interpretation of weight vectors of linear models in multivariate neuroimaging. *NeuroImage*, *87*(November), 96–110. https://doi.org/10.1016/j.neuroimage.2013.10.067

van Son, D., De Blasio, F. M., Fogarty, J. S., Angelidis, A., Barry, R. J., & Putman, P. (2019). Frontal EEG theta/beta ratio during mind wandering episodes. *Biological Psychology*, *140*(November 2018), 19–27. https://doi.org/10.1016/j.biopsycho.2018.11.003
